# Dual IgA/IgG family autoantibodies from individuals at-risk for rheumatoid arthritis identify an arthritogenic strain of *Subdoligranulum*

**DOI:** 10.1101/2022.06.09.495381

**Authors:** Meagan Chriswell, Adam R. Lefferts, Michael Clay, Alex Hsu, Jennifer Seifert, Marie L. Feser, Cliff Rims, Michelle Bloom, Elizabeth A. Bemis, M. Kristen Demoruelle, Kevin D. Deane, Eddie A. James, Jane H. Buckner, William H. Robinson, V. Michael Holers, Kristine A. Kuhn

## Abstract

The mucosal origins hypothesis of rheumatoid arthritis (RA) proposes a central role for mucosal immune responses in the initiation and/or perpetuation of the systemic autoimmunity that occurs with disease. However, the connection between the mucosa and systemic autoimmunity in RA remains unclear. Using dual IgA/IgG family plasmablast-derived monoclonal autoantibodies obtained from peripheral blood of individuals at-risk for RA, we identified cross-reactivity between RA-relevant autoantigens and bacterial taxa in the closely related families *Lachnospiraceae* and *Ruminococcaceae*. After generating bacterial isolates within *Lachnospiraceae/Ruminococcaceae* genus *Subdoligranulum* from the feces of an individual, we confirmed monoclonal antibody binding as well as MHC class II dependent CD4+ T cell activation in RA cases compared to controls. Additionally, when *Subdoligranulum* isolate 7 but not isolate 1 colonized germ-free mice, it stimulated Th17 cell expansion, serum RA-relevant IgG autoantibodies, and joint swelling reminiscent of early RA with histopathology characterized by antibody deposition and complement activation. Systemic immune responses were likely due to the generation of colon isolated lymphoid follicles (ILFs) driving increased fecal and serum IgA by isolate 7, as B and T cell depletion not only halted intestinal immune responses but also eliminated detectable clinical disease. In aggregate, these findings demonstrate one mucosal mechanism in RA through which an intestinal strain of bacteria can drive systemic autoantibody generation and joint-centered antibody deposition and immune activation.

**One-sentence summary:** *Subdoligranulum spp.* targeted by rheumatoid arthritis (RA)-relevant autoantibodies activate T cells in individuals with RA, and in mice stimulate autoantibodies and joint swelling associated with antibody deposition and complement activation.

## Introduction

The natural history of rheumatoid arthritis (RA) has been the focus of study for years, yet the causal triggers of RA remain unclear. Biomarkers that provide insights into disease mechanisms include two types of autoantibodies – anti-citrullinated protein antibodies (ACPA) and rheumatoid factor (RF) – that develop years before disease onset and are predictive for future joint disease onset and severity(*5–8*). During this preclinical period of time, individuals are considered “at-risk” for RA(*9*).

ACPA preferentially bind citrulline-containing epitopes on an array of proteins, and can contribute to joint damage in a murine model of RA(*10*). RF targets the constant region of immunoglobulin, and persistently elevated titers correlate with clinical disease activity(*11*). Notably, these antibodies arise from separate B cell lineages and mechanisms(*12*), suggesting that they function differentially in the development of RA. ACPA+ B cells undergo upregulation of genes that promote T-cell dependent responses, while RF+ B cells upregulate transcriptional programs that invoke innate immunological memory reactivation(*12*).

With regard to the initial development of RA-related autoantibodies, the mucosal origins hypothesis suggests that environmental interactions and chronic inflammation at mucosal surfaces may be important early drivers of RA pathogenesis(*13*). In support of this paradigm, ACPA are detected in the lung of individuals with longstanding RA(*14*), as well as in the sputum of individuals at-risk for RA(*15*), and this is associated with the presence of elevated cytokines, complement activation and neutrophil extracellular trap (NET) formation(*16*). The oral mucosa is also implicated, as supported by the finding that *Porphyromonas gingivalis,* a bacteria causally linked to chronic periodontitis, encodes an enzyme capable of citrullinating proteins(*17*), and ACPA+ individuals exhibit a higher relative abundance of *P. gingivalis* in their oral microbiome(*18*). Another potential oral pathway links *Aggregatibacter actinomycetemcomitans* that are expanded in the periodontium of individuals with chronic RA(*19*), and have the capacity to induce host cell hyper-citrullination(*20*) rather than direct enzymatic citrullination of proteins. Furthermore, an orally-derived strain of *Streptococcus* has been found to be capable of inducing arthritis in SKG mice(*28*). Additional data from both human and murine studies implicate the intestinal mucosa. Expansion of *Prevotella copri* in the gut of individuals with new-onset RA(*21*) and during the preclinical phase(*22*) has been described, while other groups demonstrate perturbations in *Lactobacilli*(*23*) and rare lineages of bacteria(*24*) in RA that can modulate experimental arthritis models. A further relationship to *Prevotella* is suggested by the finding that HLA-DR-containing peptides, eluted from antigen-presenting cells of patients with chronic RA, derived from these bacteria share homology with two novel RA-specific autoantigens(*25–27*). Thus, certain lines of evidence support immune responses to bacteria at mucosal surfaces as modifiers of RA relevant autoantibodies and amplifiers of underlying joint inflammation. However, despite these clinical associations, none of the proposed bacterial lineages have demonstrated an arthritogenic effect in wild type mice in the absence of other strongly pro-inflammatory factors.

Plasmablasts are a potentially highly informative subset of B cells, as they are circulating components of ongoing local immune responses(*29–31*) and could link immune responses at the mucosa with the joint. Recently, Kinslow et al. described a significant expansion of IgA+ plasmablasts in the circulation of individuals at-risk for RA(*32*). Following variable region sequencing, a substantial number of the isolated plasmablasts were found to arise in shared clonal families with both IgA+ and IgG+ members. These findings suggested a shared mucosal and systemic immune response evolution, although the specific triggers and mechanisms by which this conversion might happen remains unknown. We hypothesized that the antibodies produced by these dual IgA/IgG-containing family plasmablasts would recognize a mucosa-associated bacterium that could stimulate the development of autoimmunity. In this report, we utilized informative monoclonal antibodies (mAbs) derived from circulating dual IgA/IgG-containing plasmablast families to identify candidate intestinal bacteria in feces from at-risk individuals. From that source, we isolated a novel bacterial strain that is targeted by both dual IgA/IgG family plasmablasts and CD4+ T cells from RA patients. We show that this bacterial strain stimulates RA-related autoantibodies and joint swelling with both IgG/IgA deposition and complement activation, and demonstrate a B- and T-cell dependent pathway by which these pathologic changes arise. This novel human bacterial strain, classified as a *Subdoligranulum didolesgii*, provides a causal link between the mucosal immune system, RA autoimmunity and joint-centered pathology. Studies of this strain will provide a clinically relevant model for understanding of the drivers of microbially-driven RA-related autoimmunity.

## Results

### A subset of dual IgA/IgG family plasmablast-derived monoclonal antibodies target RA-relevant antigens as well as bacteria in families *Lachnospiraceae* and *Ruminococcaceae*

We previously identified a population of circulating plasmablasts that belong to dual IgA/IgG clonal families in individuals at risk for RA(*32*). Hypothesizing that these plamablasts may inform a mucosal to systemic immune response conversion leading to targeting of RA-relevant antigens, we established mAbs derived from plasmablasts (PB-mAbs) from shared IgA/IgG clonal families. Plasmablasts were isolated from individuals at-risk for the development of RA (n=4) and individuals with early RA (<1 year from diagnosis; n=2; Supp. Tables 1,2), and mAbs were selected for further study due to their ability to bind RA-relevant citrullinated antigens(*32*). Sequences from the variable regions were cloned onto a mouse IgG2a framework and expressed. A total of 94 successfully generated PB-mAbs confirmed binding of numerous RA-relevant autoantigens by multiplex array (Fig 1A, Supp. Table 3).

**Figure 1:**
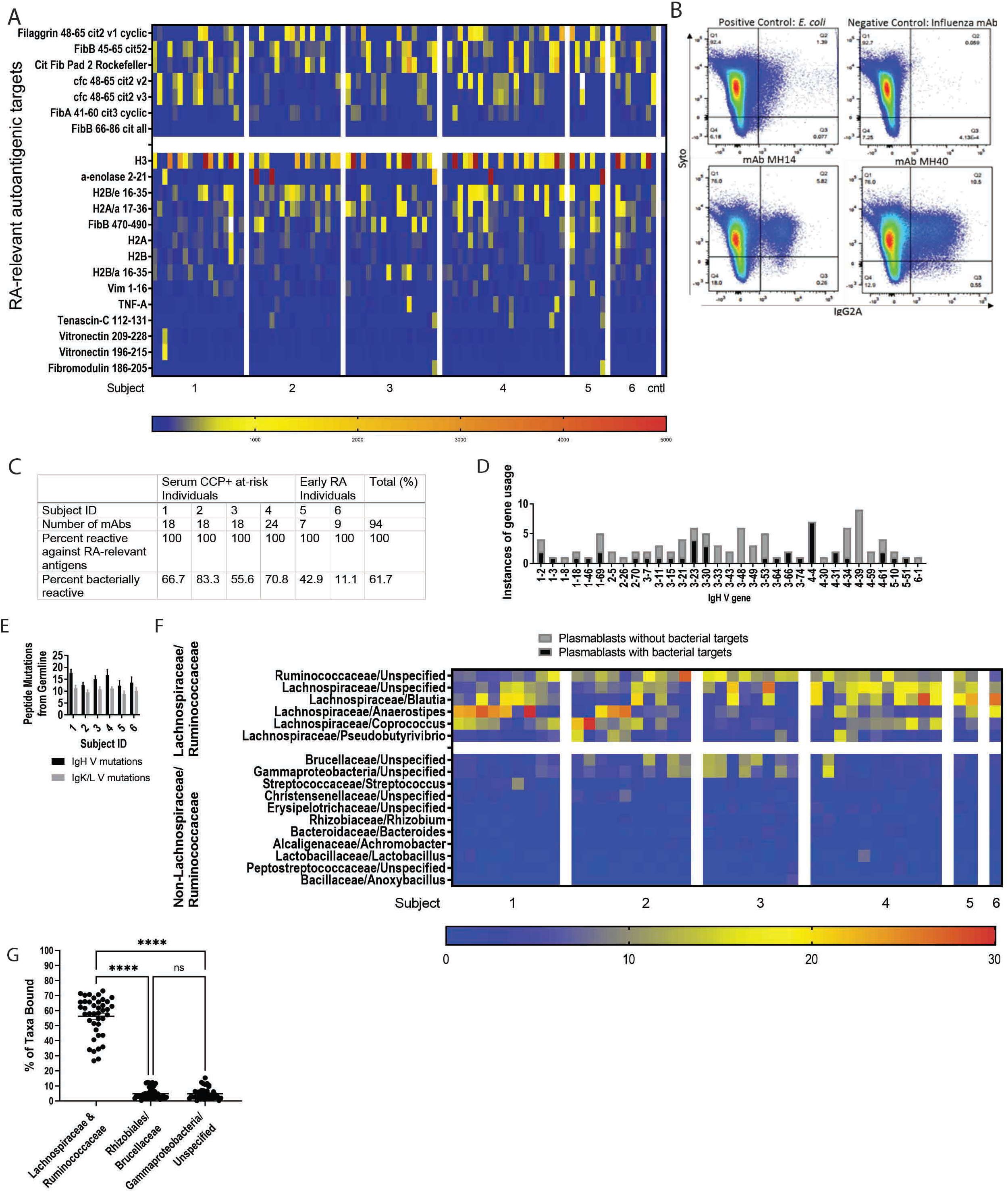
A subset of dual IgA/IgG family plasmablast-derived monoclonal antibodies cross-react with RA-relevant antigens and predominantly bind families *Lachnospiraceae* and *Ruminococcaceae.* (**A**) 94 plasmablast-derived mAbs from at-risk (n=4) and eRA (n=2) individuals belonging to shared IgG and IgA clonal families were applied to a planar array containing 346 different citrullinated and native peptide targets. The heatmap demonstrates degree of reactivity between individual PB-mAbs (columns, x-axis) with specific antigens (rows, y-axis). (**B, C**) These plasmablast-derived mAbs were screened for cross-reactivity against a broadly representative pool of fecal bacteria created from feces from individuals with eRA (n=5), at-risk (n=8) and healthy controls (n=5). The fecal pool was exposed to each antibody and analyzed by flow cytometry. The samples that were >2SD above background staining were considered to be positive for bacterial binding. Representative flow plots demonstrating binding to Syto9Green (a nucleic acid stain, x-axis) and PE (representing mAb binding, y-axis) are shown. The top left displays binding to *E. coli* (positive control), the top right to an influenza mAb (negative control). The bottom two graphs are representative binding plots of two of the positive mAbs (14 and 40). (C) The table summarizes PB-mAb reactivity against RA-relevant antigens and bacteria, separated by subject. (**D**) PB-mAbs with and without bacterial targets were analyzed for Vh gene usage. The count of mAbs expressing each Vh gene is displayed (y-axis) against represented Vh gene segments (x-axis). mAbs with bacterial targets are displayed in grey and mAbs without bacterial targets are displayed in black. (**E**) The amino acid peptide mutations from germline (y-axis) are demonstrated for each human subject (x-axis). IgH V mutations are displayed in black and IgL/K mutations are displayed in grey. (**F**) The mAb bound bacterial fraction underwent 16S rRNA sequencing and taxonomic identification. The heatmap displays percentage of total bacteria bound represented by each taxa displayed; the top taxa bound for all mAbs are shown. Each mAb is shown (x-axis), segregated by the individual from whom it was derived. Bacteria are segregated by either belonging to families *Lachnospiraceae* or *Ruminococcaceae* (top of y-axis) or not belonging to either *Lachnospiraceae* or *Ruminococcaceae* (bottom of right axis). (**G**) The three most bound taxa from each PB-mAb are represented out of the total bacteria bound (x-axis). The percent of total bacteria bound is quantified for each mAb (y-axis) and shown as round symbols for the individual mAbs and mean ± SEM as bars. P<0.0001 by one-way ANOVA with Tukey’s post-test.

Because the PB-mAbs and many circulating autoantibodies include the mucosal IgA isotype, we queried if the PB-mAbs could target intestinal bacteria. For broad representation of fecal bacteria, we pooled feces from healthy individuals (n=5), individuals at-risk for RA (n=8), and individuals with early RA (n=5; Supp. Table 4, Supp. Fig. 1A). By 16S rRNA sequencing we verified high bacterial diversity in this pool (Shannon-H Index Value 2.524; Supp. Fig. 1B. Using a negative control PB-mAb targeting influenza and a positive control mAb to *E. coli*, we developed a flow cytometry assay to identify PB-mAb binding to fecal bacteria (Fig 1B). Staining greater than two standard deviations above the mean fluorescence intensity of the negative controls was considered positive binding. Using this cut-off value, a total of 61.7% (58/94) of the PB-mAbs targeted intestinal bacteria (Fig 1C), although there was no association between isotype (IgA vs IgG) and bacterial binding (Supp. Fig 2A). There was no association between autoantigen binding preference and binding of bacterial targets (Supp. Fig 2B). However, mAbs that bound bacteria utilized a sub-set of Vh genes compared to the non-bacteria binding mAbs (Fig 1D, Supp. Fig 2C). Additionally, the PB-mAbs were substantially mutated from germline, although the number of mutations in the bacteria binding versus non-binding PB-mAbs did not significantly differ, indicating that these plasmablasts are likely not from a natural antibody pool (Fig 1E, Supp. Fig 2D).

To define the bacterial targets of the PB-mAbs, we flow sorted the mAb-bound bacterial fraction and sequenced the bacterial 16S rRNA. Predominantly targeted taxa included the closely related families *Lachnospiraceae* and *Ruminococcaceae* (Fig 1F and Supp. Fig 3A). Indeed, >50% of the total bacteria bound by the bacteria-binding mAbs combined were from families *Lachnospiraceae* and *Ruminococcaceae* (Fig 1G). These data demonstrate an interesting antibody cross-reactivity between bacterial targets, specifically *Lachnospiraceae* and *Ruminococcaceae,* among a subset of mAbs that bind RA-relevant autoantigenic targets.

### *Ruminococcaceae Subdoligranulum* strains isolated from a human sample are targeted by plasmablast mAbs and stimulate CD4+ T cells from RA patients in a DR-dependent manner

In order to investigate immune responses to *Lachnospiraceae* and *Ruminococcaceae* in RA, we first established primary bacterial isolates from an individual with >40% abundance of these taxa in their feces (Supp. Fig 3B). A total of 50 isolates were established, seven identified as *Lachnospiraceae/Ruminococcaceae* by qPCR (isolates 1-7), and five confirmed as pure isolates by 16S rRNA sequencing (isolates 1, 3-5, and 7). These five isolates underwent whole genome sequencing.

Sequences were assembled, scaffolded, and cleaned using Abyss(*33*), input into the Biobakery Workflow(*34*), and categorized as unidentified species within *Ruminococcaceae Subdoligranulum* (Fig. 2A), which we designate as *didolesgii*. We narrowed down the five isolates to two candidates (isolates 1 and 7) for further study due to their differing abilities to induce joint swelling in monocolonized mice (Supp. Fig. 5 and elaborated below in Fig. 3). We confirmed that a subset of PB-mAbs (numbers 4, 28, 58, and 91), selected based on their binding of both *Lachnospiraceae* and *Ruminococcaceae* in Fig 1, bound our bacterial isolates, while mAb 7 that did not bind *Lachnospiraceae* or *Ruminococcaceae* did not bind isolates 1 or 7 (Fig 2B).

**Figure 2:**
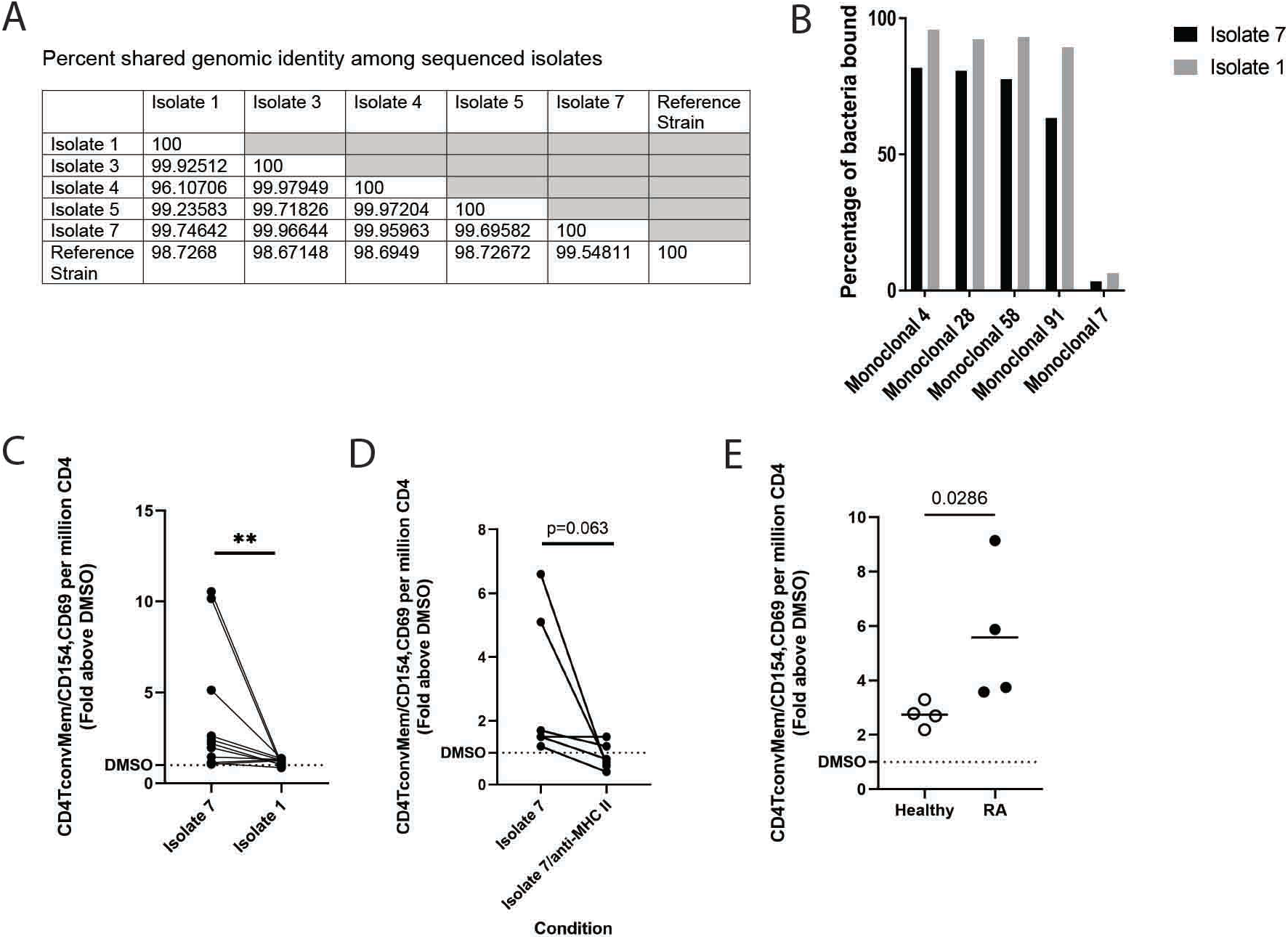
*Ruminococcaceae Subdoligranulum* strains isolated from a human sample are targeted by plasmablast mAbs and also stimulate CD4+ T cells from RA patients in a DR-dependent manner. (**A**) Seven primary strains of *Ruminococcaceae Subdoligranulum* were isolated from the feces of an individual. 5 isolates were selected for short read genome sequencing based on taxonomic identification by 16S rRNA sequencing. The table represents percent genomic shared identity among the 5 isolates as well as against a reference genome found to be genetically aligned (MGYG-HGUT02424; unidentified genus in Order *Clostridiales*, which includes *Lachnospiraceae* and *Ruminococcaceae*). (**B**) Isolates 1 and 7 were matched against four selected PB-mAbs (numbers 4, 28, 58, and 91) that bound highly to other various patterns of *Ruminococcaceae* and *Lachnospiraceae* species to verify that they targeted the strains. They were also matched against a control mAb (number 7) that was previously found to not bind bacteria. The percent of bacteria bound to mAb is displayed (y-axis) against each selected mAb (x-axis). Binding by isolate 7 is shown in black, and binding to isolate 1 is shown in grey. (**C**) Human PBMCs from RA cases (n=11) were stimulated with isolate 1 or isolate 7 and evaluated by flow cytometry for T cell activation as measured by presence of CD154 and CD69. Fold change of the CD4 T cell activation response relative to DMSO control (y-axis) is displayed for isolates 1 and 7 (x-axis). Each individual is represented by a round symbol with a line connecting the paired samples. **,P<0.01, non-parametric Wilcoxon matched-pairs signed rank test. (**D**) A Class II HLADR (clone L243) block was applied prior to stimulation. Fold change of the CD4 T cell activation as measured by CD154 and CD69 relative to DMSO control (y-axis) is displayed. Each individual is represented by a round symbol with a line connecting the paired samples. P=0.625, non-parametric Wilcoxon matched-pairs signed rank test. (**E)** Isolate 7 specific responses among T cells was tested, comparing cells derived from RA cases to healthy control T cells. Fold change of the CD4 T cell activation as determined by CD154 and CD69 relative to DMSO control (y-axis) is displayed, comparing healthy control cells to RA case cells (x-axis). Each round symbol represents an individual and the bars as mean for the group. P=0.0286, Mann-Whitney test.

**Figure 3:**
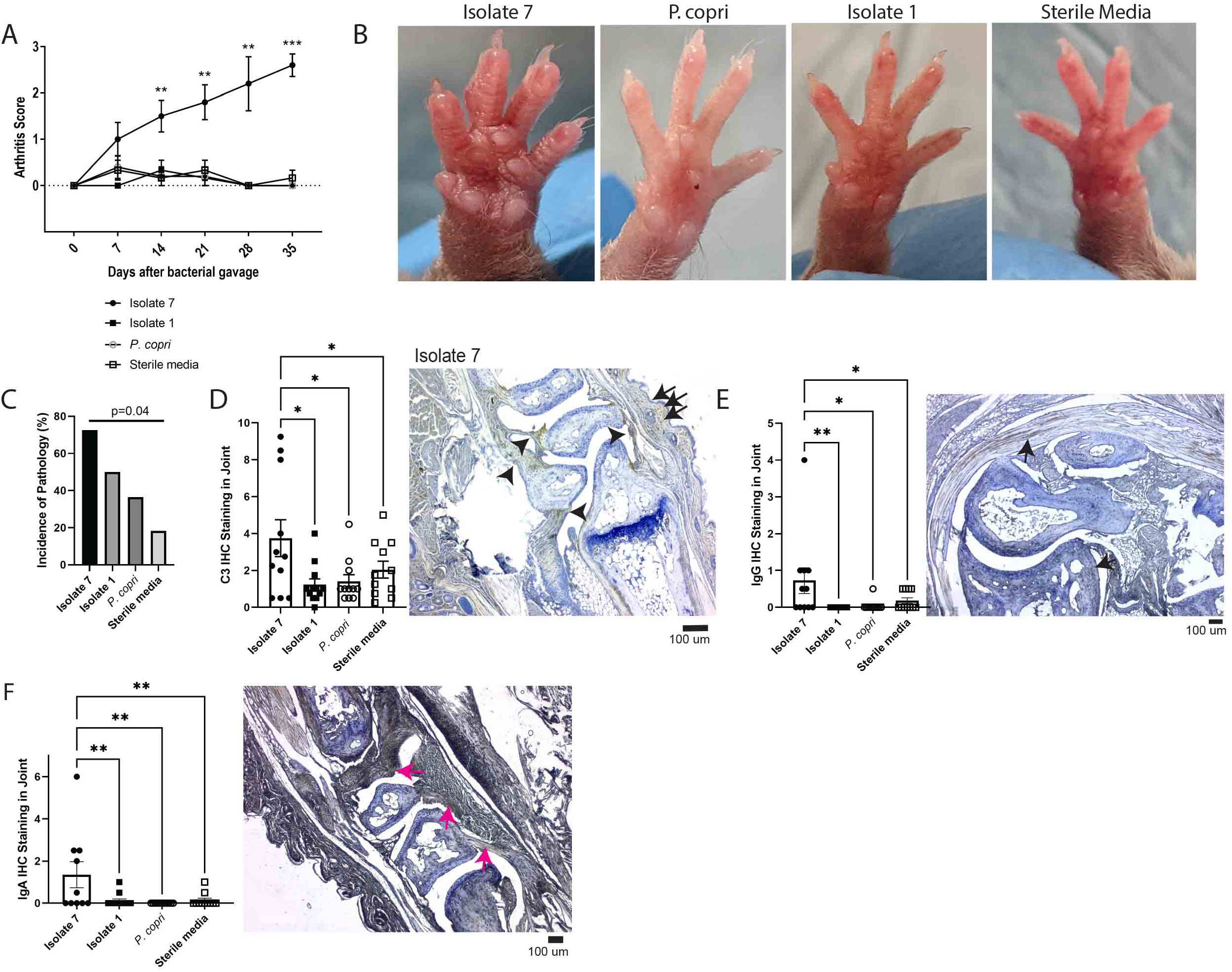
*A specific Subdoligranulum* strain stimulates joint swelling and inflammation in mono-colonized mice that is characterized by IgG, IgA, and complement C3 deposition in joints. (**A and B**) *Subdoligranulum* isolates 1 and 7 as well as *Prevotella copri* or sterile media were gavaged separately into germ-free DBA/1 mice (n=6 isolate 1, n=6 isolate 7, n=5 *P. copri*, and n=6 sterile media). Mice were observed weekly for 35 days for the development of joint swelling and assessed a score based on the number of joints affected. (**A**)The mean ± SEM score is shown (y-axis) over time after bacterial gavage (x-axis). **,P<0.01and ***,P<0.001, unpaired t-test. (**B**) Representative photographs of paws from the treatment groups are shown to demonstrate the swelling observed in mice monoc-colonized with isolate 7. (**C**) Paw histology was assessed by a pathologist in a blinded fashion. Displayed is the incidence of pathology (y-axis) separated by treatment group (x-axis). P=0.04, Fisher’s exact test. (**D**) IHC for the C3 component of complement was performed on decalcified paw sections. C3 staining intensity scores are displayed (y-axis) separated by treatment group (x-axis). Symbols represent individual mice while bars are the mean ± SEM. *,P<0.05, one-way ANOVA with Bonferroni post-test. A representative section for isolate 7 is displayed, with deposition indicated by arrows; the scale bar represents 100µm. (**E**) IgG deposition in decalcified paw sections was assessed by IHC. Intensity scores with symbols as individual mice and bars as mean ± SEM are displayed (y-axis), separated by treatment group (x-axis). *,P<=0.05 and **P<0.01, one-way ANOVA with Tukey post-test. A representative section is displayed, with deposition indicated by arrows; the scale bar represents 100µm. (**F**) IgA deposition in decalcified paw sections was assessed by IHC and intensity scores are displayed (y-axis), separated by treatment group (x-axis). Symbols represent individual mice while bars are the mean ± SEM. **P<=0.01, one-way ANOVA with Tukey post-test. A representative section is displayed, with deposition indicated by arrows; the scale bar represents 100µm. (n=12 isolate 1 gavaged, n=11 isolate 7 gavaged, n=11 *P. copri* gavaged, and n=11 sterile media gavaged, across two experiments).

Next, we assessed if the *Subdoligranulum* isolates were recognized by circulating T cells from individuals with RA to further support their immunologic relevance. PBMCs from 11 individuals with RA (Supp. Table 5) were stimulated with 50 ng/ml oxygen-killed *Subdoligranulum* isolate 1 or isolate 7 for 14 hours. Compared to isolate 1, isolate 7 significantly activated CD4+ T cells in the PBMCs as measured by accumulated surface CD69 and CD154 expression (Fig. 2C, Supp. Fig. 4). T cell activation was MHC class II dependent as blocking with anti-HLADR4 abrogated CD69 and CD154 expression (Fig. 2D). Furthermore, another experiment indicated that isolate 7 specific responses were more prevalent in T cells derived from RA cases as compared to healthy controls (Fig 2E, Supp. Table 6). Our findings support the hypothesis that the *Subdoligranulum* isolate 7 is immunologically relevant and suggest that strain variations may be of importance to understanding the pathogenesis of RA.

### *Subdoligranulum* isolates stimulate joint swelling characterized by antibody and complement deposition in mono-colonized mice

Having associated immunity towards *Subdoligranulum* isolates with RA and deriving them from a human sample we then investigated if the isolates were capable of inducing RA-relevant autoimmunity in mice. *Subdoligranulum* isolates 1, 3-5, and 7 were gavaged at 5 x 10^6^ CFU into germ-free DBA/1 mice. Given the previously published association of *Prevotalla copri* with new-onset RA(*21*) and the homology between proteins in this bacteria and RA-relevant autoantigens(*25–27*), we mono-colonized a group of germ-free DBA/1 mice with *P. copri* (DSMZ 18205(*35*)) as a control in addition to a sterile media gavage. Four of the five strains, and neither control, caused joint swelling starting ∼14 days after gavage and persisted until 35 days when most mice were euthanized (Fig 3A,B and Supp. Fig. 5A), although when followed through 63 days, arthritis was still observed (Supp. Fig. 5B). Interestingly, germ-free C57Bl/6J mice also developed very mild but detectable joint swelling after mono-colonization with isolate 7 (Supp. Fig. 5C).We confirmed equal, stable bacterial colonization across groups (Supp. Fig. 5D). The finding of arthritis development with introduction of a single strain alone was unexpected, as non-transgenic murine models of joint disease typically require intradermal injection of antigen with adjuvant, adjuvant alone, or intravenous transfer of pathogenic antibodies to develop disease(*36, 37*).

Evaluation of pathology demonstrated a range of synovitis, osteitis, vasculitis, and soft tissue inflammation, all mild in severity (Supp. Fig. 6). Yet all-combined, mice colonized with isolate 7 had the highest incidence of pathology compared to the other treatment groups (p=0.04, Fig. 3C). Despite mild immune cell infiltrate, immunohistochemistry of the joints for immunoglobulin deposition and complement protein C3 demonstrated marked deposition both in the joint space and intradermally (Fig 3D, Supp. Fig 7A). Additionally, immunohistochemistry demonstrated IgG and IgA deposition in the joint (Fig 3E,F, Supp. Fig 7B,C). These findings suggest that an antibody mediated process may be a key factor in driving the joint swelling in this murine phenotype.

### Serum IgA, RA-related autoantibodies, and splenic Th17 cells expand in *Subdoligranulum* isolate 7 mono-colonized mice

In order to understand the systemic immune response following gavage with isolate 1, isolate 7, *P. copri*, or sterile media, serum from mice was collected on days 14 and 35 after gavage. The total serum IgA was significantly increased in isolate 7 mono-colonized mice at day 14 as compared to the other groups (Fig. 4A), but this normalized by day 35 (Supp. Fig. 8A) and was not observed for IgG (Supp. Fig. 8B).

**Figure 4:**
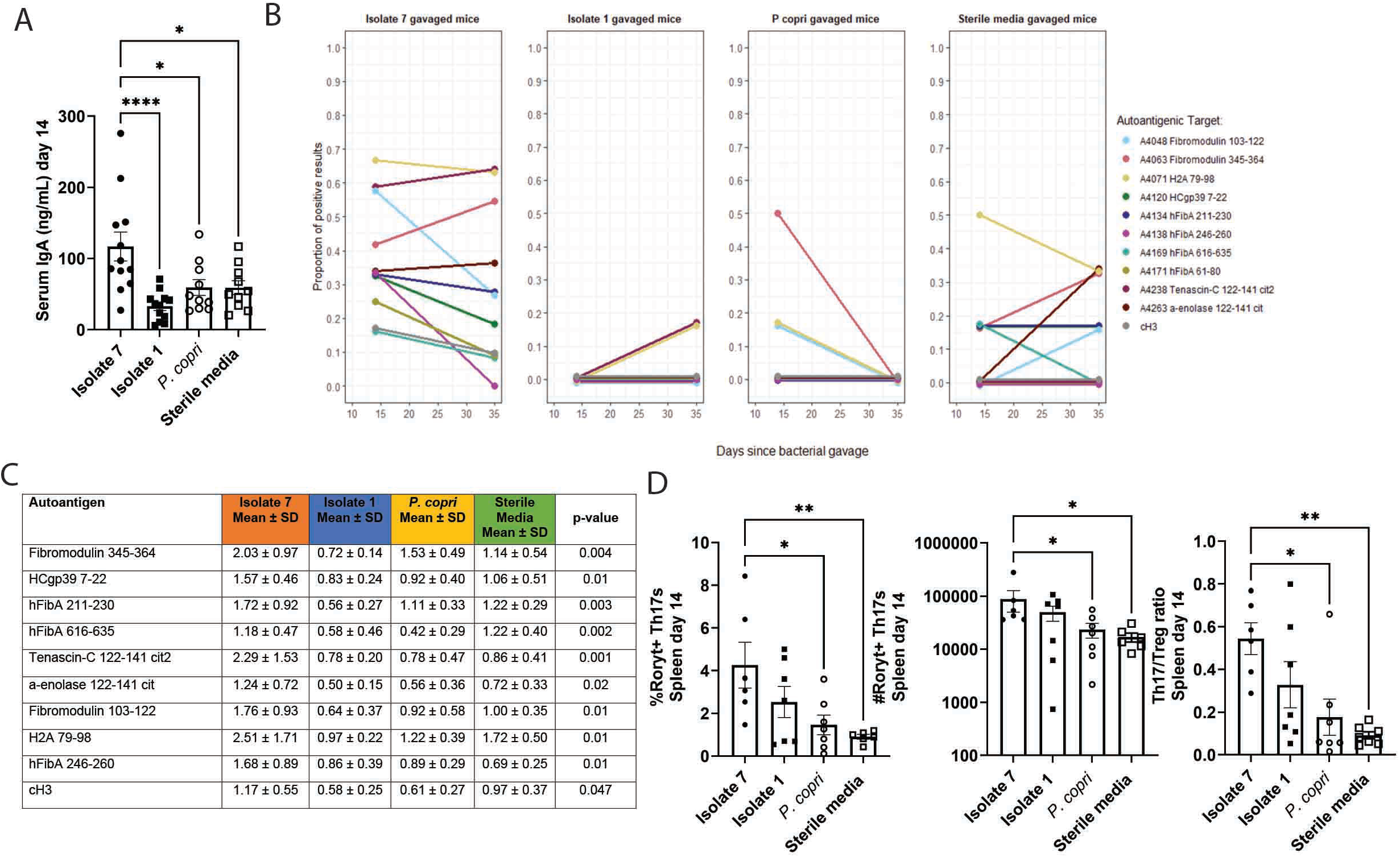
*Subdoligranulum* isolate 7 causes development of increased serum IgA, systemic RA-related autoantibodies, and expanded splenic Th17 populations. (**A-C**) Serum from mice mono-colonized with either isolate 1 (n=6), isolate 7 (n=12), or *P. copri* (n=6), or given sterile media (n=6) was collected at days 14 and 35 after gavage. (**A**) The total serum IgA at 14 days after gavage was determined by ELISA; serum IgA is displayed (y-axis) against treatment group (x-axis). Symbols represent individual mice while bars are the mean ± SEM. *,P<0.05 and ****,P<0.0001, Kruskal-Wallis test with Dunn’s post-test. (**B**) Sera was analyzed on a planar array containing ∼350 citrullinated and native peptides for autoantigens relevant in RA. A cutoff for positivity was established at the 80^th^ percentile of autoantibody reactivity for individual samples on this assay (1.5 relative units) and the proportion of murine samples meeting or exceeding this threshold (11 antigens as displayed by) at each timepoint is shown (y-axis). Each treatment group is shown from left to right. (**C**) Mean ± SD values for serum reactivity from each treatment group with specific RA-relevant autoantigens shown in panel (B) are shown in the table, comparing the four groups. P-values were determined by Kruskal-Wallis test with Dunn’s post-test. (**D**) Spleens were collected from each mouse at 35 days post gavage and CD4+ T cell populations were analyzed by flow cytometry (n=7 isolate 7, n=7 isolate 1, n=6 *P. copri*, n=8 sterile media). The percentage (left) and absolute number (middle) of Roryt+ Th17 cells as well as the Th17 to Treg ratio (right) is displayed. Symbols represent individual mice while bars are the mean ± SEM. *,P<0.05 and **,P<0.01 using Kruskal-Wallis with Dunn’s post-test.

We next evaluated serum autoantibodies using a planar array containing ∼350 RA-relevant autoantigens. Isolate 7 gavaged mice developed and maintained serum autoantibodies against RA-relevant autoantigens at higher proportions 14-35 days after gavage (P<0.05) than mice in the other groups (Fig 4B). Furthermore, isolate 7 mono-colonized mice developed several autoantibodies targeting RA-relevant antigens such as fibromodulin at greater titers in comparison to the other groups (Fig. 4C). Notably, there was reactivity to both citrullinated and native peptides, analogous to the reactivity profiles of the human PB-mAbs (Fig. 1A). These data indicate that mono-colonization with isolate 7 allows for the establishment of persistent RA-relevant autoantibodies in circulation.

Given CD4+ T cell reactivity to isolate 7 in patients with RA, we evaluated splenic T cell populations at days 14 and 35 in mice mono-colonized with isolate 1, isolate 7, or *P. copri*, or gavaged with sterile media (Supp. Fig 9A). We focused on Th17 and Treg populations due to the demonstrated role of intestinal microbiota in their development(*39, 40*), and on Tfh because of our observed changes in autoantibodies. At day 14, splenic Th17 cells were significantly increased in percentage, absolute number, and Th17/Treg ratio in isolate 7 mono-colonized mice compared to *P. copri* and sterile media gavaged mice (Fig 4D, Supp. Fig. 9B, C). However, we did not observe significant differences in Treg or Tfh subsets in the isolate 7 group (Supp. Fig. 9B, C). By day 35, the expansion of Th17 cells remained in isolate 7 mono-colonized mice compared to sterile media gavaged mice (Supp. Fig. 10). Our data are consistent with others’ findings of Th17 expansion aiding in the development of autoantibody-mediated arthritis in mice(*41*) and in the evolution of human RA(*42, 43*). These findings in aggregate suggest a possible mucosal to systemic immune system response driven by IgA, resulting in the targeting of autoantigens by the immunoglobulins that reach the systemic circulation after being educated in the mucosa.

### *Subdoligranulum* isolate 7 generates intestinal isolated lymphoid follicles, increased IgA, and Th17 skewing

We next investigated the effects of *Subdoligranulum* isolate 7 on intestinal immunity. First, to determine if isolate 7 relative to isolate 1, *P. copri*, or sterile media affected intestinal permeability, FITC-dextran was administered orally to mice four hours prior to euthanasia, at which time sera was collected and the levels of FITC-dextran measured. All three mono-colonizations resulted in improved barrier compared to sterile media (Fig 5A), indicating that the two *Subdoligranulum* isolates and *P. copri* were capable of at least partially restoring the barrier defect of germ-free mice(*44*). Intestinal histology revealed significantly increased numbers of isolated lymphoid follicles (ILFs) in isolate 7 colonized mice as compared to *P. copri* and isolate 1 colonized mice, and a trend towards increased ILF size in isolate 7 gavaged mice (p=0.09 compared to sterile media;Fig 5B, C), without change of colonic crypt depth or small intestinal villus morphology among isolate 7 gavaged mice (Supp. Fig 11). Interestingly, there is an increase in villus width and crypt depth among isolate 1 gavaged mice which could suggest a mild injurious intestinal effect among mice gavaged with this strain (Supp. Fig 11). The ILFs in isolate 7 colonized mice were larger and more numerous as compared with other groups, and more closely resembled classical mature ILF morphology(*45*) (Fig 5C), suggesting increased mucosal IgA generation(*46*). Indeed, luminal IgA secretion was significantly increased in isolate 7 colonized mice as compared to the other treatment groups (Fig 5D), although similar to the serum, this difference resolved by day 35 (Supp. Fig 12A). There was no significant difference in fecal IgG among groups (Supp. Fig 12B). In associated mucosal lymphoid tissues mesenteric lymph nodes (MLNs) and Peyer’s patches (PPs), the ratio of Th17/Treg T cell subsets increased significantly in the isolate 7 mono-colonized mice compared to sterile media gavaged mice at day 14 following gavage (Fig 5E), although the percentages and absolute numbers were not significantly different (Supp. Fig 13 A-C). In aggregate, our observations suggest that *Subdoligranulum* isolate 7 stimulates a robust intestinal immune response characterized by the formation of ILFs functioning to secrete IgA.

**Figure 5:**
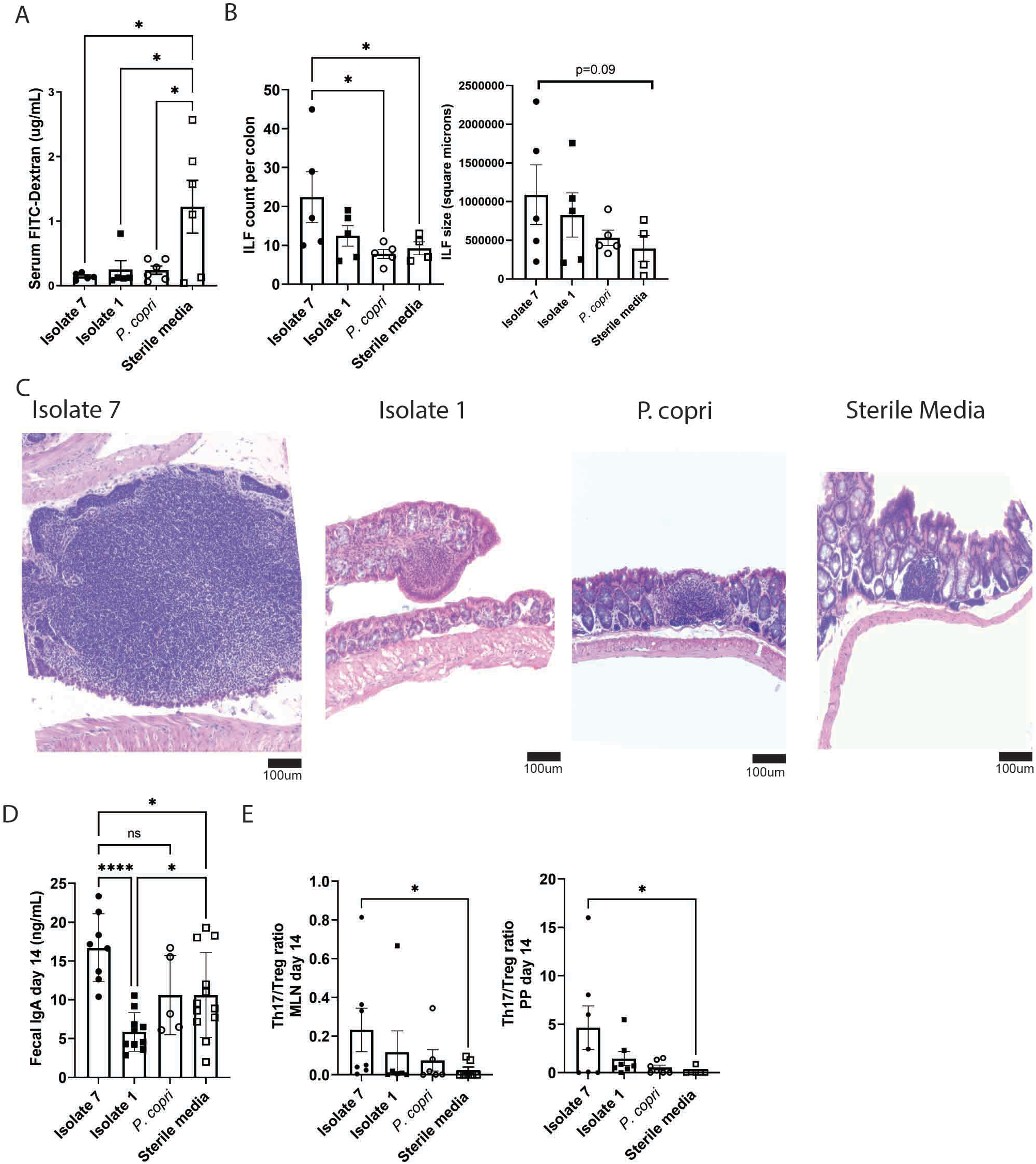
*Subdoligranulum* isolate 7 causes development of intestinal isolated lymphoid follicles, increased mucosal IgA and Th17 skewing in mucosal lymphoid tissues. (**A**) FITC-Dextran was orally gavaged into mice mono-colonized with isolate 1 (n=5), isolate 7 (n=7), *P. copri* (n=6) or sterile media (n=6) 4 hours before euthanasia. At the time of euthanasia, serum was collected and tested for the presence of FITC-Dextran. Concentration of serum FITC-dextran is displayed (y=axis) against each treatment group (x-axis). Symbols represent individual mice while bars are the mean ± SEM.*,P<0.05, one-way ANOVA with Tukey post-test. (**B and C**) Colon histology from each group was analyzed. (**B**) The number (left) and size in area (right) of isolated lymphoid follicles (ILFs) per colon (y-axis) separated by treatment group (x-axis) is shown. Symbols represent individual mice while bars are the mean ± SEM. *,P<0.05, one-way ANOVA with Tukey post-test. (**C**) Representative colon histology containing ILFs in each group are shown. Scale bars represent 100µm. (**D**) Feces from mono-colonized mice at day 14 after gavage were tested for total IgA by ELISA (n=11 isolate 7 gavaged, n=12 isolate 1 gavaged, n=6 *P. copri* gavaged, n=12 sterile media gavaged). Symbols represent individual mice while bars are the mean ± SEM. *,P<0.05; ****P<0.0001; and ns=not-signficant by Kruskal-Wallis with Dunn’s post-test. (**E**) Mesenteric lymph nodes (MLNs) and Peyer’s patches (PPs) were evaluated at 14 days post-gavage for Th17 and Treg subpopulations by flow cytometry (n=7 isolate 7 gavaged, n=7 isolate 1 gavaged, n=7 *P. copri* gavaged, n=8 sterile media gavaged). The ratio of Th17/Treg cells is displayed with individual mice shown as symbols while bars are the mean ± SEM.. *P<0.05by Kruskal-Wallis with Dunn’s post-test.

### Joint swelling in *Subdoligranulum* isolate 7 colonized mice is dependent on T and B cells but not granulocytes

To determine if the observed paw swelling was truly mediated by adaptive immunity, we selectively depleted B cells, T cells, or granulocytes using mAbs (Supp. Fig 14) two days prior to mono-colonizing mice with isolate 7. The depletions were efficacious at depleting the B cells by 2-fold, the T cells by 10-fold, and granulocytes by 5-fold on average. Mice depleted of T or B cells did not develop paw swelling, whereas control antibody treated mice did (Fig. 6A), indicating that adaptive immunity is required for our phenotype. Though mice depleted of granulocytes developed paw swelling equal to treatment with control antibody, the onset of swelling was delayed by about one week (Fig 6A), suggesting that while granulocytes aid in the phenotype, they are not essential. ILFs were not affected in the T cell or granulocyte depleted mice, though they are reduced in size in the B cell depleted mice (P=0.07, Fig 6B). Circulating and fecal IgA was significantly decreased in the B and T cell depleted mice at days 14 and 35 after bacterial gavage, and circulating and fecal IgG was decreased at day 35 after bacterial gavage (Fig. 6C-F). Interestingly, circulating IgG and IgA is also significantly reduced in the granulocyte depleted mice, suggesting that granulocyte-dependent Ig synthesis at the mucosal surface that spreads systemically may be important in this model.

**Figure 6:**
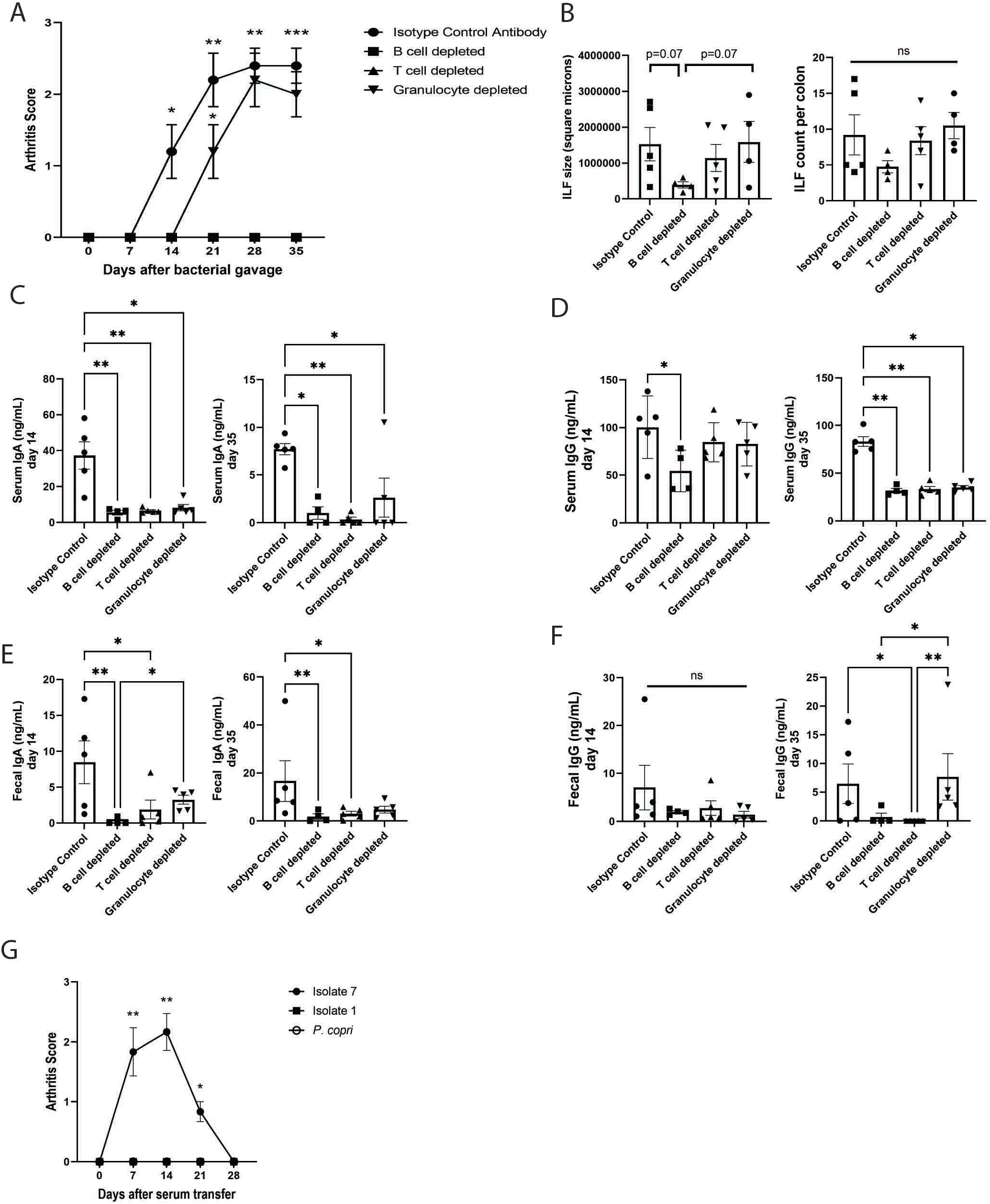
Joint swelling is dependent on T and B cells but not granulocytes. (**A**) Mice were selectively depleted or their B cells (n=4), CD4+ T cells (n=5), or granulocytes (n=5) through administration of depleting mAbs (n=5 isotype control). Joint swelling was assessed as previously and the mean mean ± SEM score (y-axis) is shown over time (x-axis). *,P<0.05 **,P<0.01 and ***,P<0.001, unpaired t-test). (**B**) ILF area in square microns (left) and numbers (right) was assessed in colon histology from isotype control and cell depleted mice at day 35 after bacterial gavage. Symbols represent individual mice while bars are the mean ± SEM. P-values were determined by one-way ANOVA with Tukey post-test. (**C-F**) Total IgG and IgA in serum and feces at days 14 and 35 post-bacterial gavage were determined by ELISA. Symbols represent individual mice while bars are the mean ± SEM. *,P<0.05; **,P<0.01; and ns=not significant as determined by Kruskal-Wallis with Dunn’s post-test. (**G**) Pooled day 35 serum from mice mono-colonized with isolate 1, isolate 7, and *P. copri* was injected into healthy germ-free DBA/1 mice, and these mice were monitored for the development of joint swelling. The mean clinical score ± SEM (n=6 isolate 7 serum transfer, n=6 isolate 1 serum transfer, n=7 *P. copri* serum transfer) is shown relative to time after-serum transfer (x-axis). P<0.05, Kruskal-Wallis with Dunn’s post-test.

As the isolate 7 colonized mice develop serum autoantibodies to RA-relevant antigens, we queried if they were pathogenic. Serum was collected from mice 35 days after colonization with isolate 1, isolate 7, or *P. copri* mono-colonized mice and pooled by colonization group. Intraperitoneal injection of serum from isolate 7 colonized mice into healthy germ-free DBA/1 mice resulted in paw swelling observed within days and similar to other serum transfer studies(*47, 48*), resolving by 28 days following transfer, whereas sera from isolate 1 and *P. copri* colonized mice were unable to stimulate this phenotype (Fig. 6H, Supp. Fig 15). In aggregate, these findings support a potential mechanism for *Subdoligranulum* isolate 7 inciting B cell autoimmunity through a local intestinal to systemic immune response.

## Discussion

The mucosal origins hypothesis for the development of RA is based on compelling human immunologic and phenotypic data: IgA plasmablasts and dual IgA/IgG clonal families are expanded in the circulation during the at-risk period preceding clinically apparent RA(*32*); ACPA have been detected at several mucosal surfaces throughout the human body(*13–15, 49*) and are often of an IgA isotype(*32*); periodontitis, *P. gingivalis*, and *A. actinomycetemcomitans* in the oral mucosa link to ACPA generation(*17, 18, 20*); and cross-reactivity between proteins in *P. copri* and RA-relevant autoantigens has been suggested(*25–27*). Nevertheless, no specific microbiota-derived organism has to date been shown to be both recognized by patients with RA and able to cause joint disease in experimental models. Other studies have demonstrated that strains can modulate existing arthritis(*28, 50, 51*), but the demonstration of the ability of a strain to incite pathology independently in genetically unmanipulated and non-mutant mice is unique. Utilizing dual IgA/IgG plasmablast derived monoclonal antibodies from human participants at risk for RA as a tool to probe the mucosal origins hypothesis, our findings establish a line of data linking a specific intestinal bacteria in the genus *Subdoligranulum* with local intestinal ILF formation, T cell and RA-related autoantibody development, and ultimately paw swelling associated with IgG, IgA, and C3 deposition in mice.

Not only do the presence of IgA plasmablasts and shared clonality with IgG plasmablasts during the at-risk period in RA suggest a mucosal trigger for ACPA generation and disease(*32*), but they also provide an important tool to probe for potential antigens driving their clonal expansion. Although the hallmark of ACPA is the citrulline specificity, recent studies of PB-derived mAbs from the peripheral blood of RA patients challenge this concept, often finding cross-reactivity with antigens containing and lacking posttranslational modifications(*52, 53*). Finding that polyautoreactive PB-mAbs from at-risk individuals similarly reacted with *Lachnospiraceae* and *Ruminococcaceae* further supports that bacteria could potentially drive a polyreactive response with RA-related antigens as suggested for ACPA. We are aware of the intricacies of the terms polyreactive and cross-reactive, and have chosen to term these PB-mAbs as cross-reactive as we hypothesize that there may be a discrete antigen(s) on the *Subdoligranulum* isolates that is a binding target for these antibodies. However, further studies are necessary to confirm the identity of this putative antigen(s) and determine whether molecular mimicry is relevant to the interaction.

The bacterium of interest in this study, that we propose naming *Subdoligranulum didolesgii* (see author’s note) is a member of a highly heterogeneous group of bacteria belonging to the family *Ruminococcaceae* that is phylogenetically interlinked with the family *Lachnospiraceae*(*54, 55*), both within order Clostridiales. As we have not yet determined whether this bacterium is a commensal organism or a pathobiont, we have avoided using these terms in this manuscript. *Ruminococcaceae* and *Lachnospiraceae* are two of the most abundant families in both the human and murine gut microbiome(*56, 57*), raising the question of why certain strains could lead to the development of arthritis without RA being a ubiquitous disease in the population. Several potential mechanisms could explain this observation. Both *Lachnospiraceae* and *Ruminococcaceae* live deep in the mucosa, occupying a niche that allows them close access to their host organism and subsequently greater immunomodulatory potential compared to bacteria localized in the lumen(*58*). Furthermore, heterogeneity in families *Lachnospiraceae/Ruminococcaceae* may be due in part to their engagement in lateral gene transfer, or alternatively infection by bacteriophages, leading to the expression of novel proteins and gain of unique functions that could lead to host immunomodulation. Interestingly, at risk individuals who are serum ACPA+ harbor increased abundances of *Lachnospiraceae* and *Ruminococcaceae* phages in their feces compared to ACPA- at risk individuals and controls(*59*). Molecular mimicry between bacterial and host proteins may lead to the development of autoimmunity, or bacterial products may alter mucosal immune system dynamics. In support of molecular mimicry is the recent discovery of T and B cell targeted autoantigens in RA with shared homology between synovial and *P. copri* proteins(*25*). Alternatively, murine models of autoimmune arthritis, as evidenced by the K/BxN, collagen-induced arthritis, SKG, and HLA-B27/β2m models, demonstrate that microbial stimulation of the Th17 pathway is required for the development of arthritis(*60–64*). Finally, community dynamics within microbial populations under different environmental pressures could lead to altered metabolite generation, which affect immune responses within the host. For example, bacterial metabolism generates short chain fatty acids like butyrate and propionate that promote Treg development and function, protecting from collagen-induced arthritis and HLA-B27/β2m-mediated arthritis(*65, 66*). Thus, *Subdoligranulum didolesgii* may be able to promote RA-relevant autoimmunity through multiple pathways.

Mice gavaged with *Sublogranulum* isolate 7 developed paw swelling in a manner that is highly reproducible through blinded scoring. While a profound immune infiltrate into the joints was absent, there was an increase in IgG, IgA, and C3 deposition. These findings could resemble stage of RA development in humans known as tenosynovitis, tendon sheath inflammation that can be associated with microscopic synovitis in the hands and feet during the at-risk and early RA periods(*38, 67*). We see evidence of mild synovial inflammation in the joints of mice colonized with *Subdoligranulum* isolate 7. If we are indeed capturing a stage of joint swelling similar to tenosynovitis in our mice, this model may be more aligned with the typical pattern of human RA development than previous murine arthritis models. Additionally, this model may be more aligned with the current two-hit hypothesis for the development of RA, the first hit being an environmental stimulus that catalyzes the development of circulating RA-relevant autoantibodies and tenosynovitis, and the second hit being an additional factor, whether genetic or environmental, that triggers the overt synovitis in classifiable RA(*13, 68*).

We were also able to appreciate substantial local histopathologic and immune changes in the gut of the mice gavaged with *Subdoligranulum* isolate 7. One key finding was the generation of mature ILFs in the colon, which are known to develop dynamically in response to bacteria in the intestine(*69*). They are comprised mostly of B cells with T cells and CD11c+ dendritic cells interspersed(*70*). Mature ILFs can form germinal centers, and the B cell repertoire inside of them has been shown to resemble systemic B cell populations(*71*). Clinically, ILF hyperplasia is linked with IgA dysfunction(*72*), and has been seen in children with circulating IgA and IgG against milk proteins(*73*), signaling the potential for mature ILFs to stimulate systemic immune responses. *Subdoligranulum* isolate 7 seems to be stimulating mature ILFs that subsequently produce an immune response that triggers a mucosal to systemic immune response conversion.

Altogether our data suggest one pathway by which the intestinal microbiome and mucosal immune responses can lead to systemic autoreactivity and joint pathology that is highly relevant to and likely a causal pathway in human RA. Numerous outstanding questions remain: What is the prevalence of our specific *Subdoligranulum* strain in the general population versus those with RA, and are there strain variations that could explain our findings? How does *Subdoligranulum* specifically interact with the host, and, what is the mechanism for T cell and B cell responses towards it? How does *Subdoligranulum* itself interact within the microbial community? And finally, what is the role of host genetics during the immune responses to *Subdoligranulum*? Nevertheless, our results highlight the utility of using host immune responses, here the IgA/IgG response in plasmablasts, in probing for relevant microbiota at mucosal surfaces and discovering a contribution to RA by a novel organism that would not be identified otherwise.

## Materials and Methods

### Study Design

The objective of this study is to understand the bacterial targets of human autoantibodies in the context of RA, to isolate these bacteria, and to understand their role as drivers of murine joint disease. For all experiments, the number of replicates, statistical test used, and *P* values are reported in the figure legends. The reported replicates refer to biological replicates, either murine or human. For human studies, individuals were recruited at either the University of Colorado or Benaroya Research Institute under IRB approval. Full details of the human cohorts utilized can be found in supplemental methods. For murine studies, mice were derived germ-free and housed at the University of Colorado gnotobiotic facility under IACUC approval. Cages of mice were randomly assigned to different treatment groups, and the assessor of joint swelling was blinded to treatment group. To ensure animal welfare, weight loss and signs of pain and distress were utilized as rationale for the premature endpoint of any study. For all histopathological analysis, the scorer was blinded to treatment group. Full experimental details can be found in the supplemental methods.

### Statistical Analysis

Unless noted elsewhere, normally distributed data as determined by D’Agostino test were evaluated by ANOVA and a post-hoc t-test with Tukey’s correction and non-parametric data were analyzed using Mann-Whitney or Kruskal Wallis with Dunn’s correction using Graphpad Prism 9 Software.

### Data Sharing

Data generated from PB-mAbs, 16S rRNA, and bacterial whole genome sequencing are publicly accessible in accordance with NIH guidelines.

## Acknowledgements

The work by the authors is supported through U01 AI101981 (VMH), Pfizer ASPIRE (KAK), T32 AR07534 (MC, ARL), R01 AR075033 (KAK), and the Rheumatology Research Foundation Future Physician Scientist Award (MC).

MC, MKD, KDD, EJ, JB, WR, VMH, and KAK designed the studies. MC, ARL, AH, JS, MF, CR, MB performed experiments and collected data. EB assisted with statistical analyses. Initial drafts of the manuscript were written by MC and KAK. All authors contributed to data analysis and interpretation as well as reviewing and revision the manuscript and approved its final form for submission.

The authors have no conflicts of interests to declare.

Sequencing data are publicly available through the NIH’s Sequence Read Archive (SRA) accession numbers SUB10802055, SUB 10768018, SUB10785477, and SUB10768012.

## Author’s Note

Didolesgi (ᎱᏳᎩ) is the Cherokee word for arthritis or rheumatism. The first author is an enrolled member of the Cherokee Nation of Oklahoma, and we chose this name for a few significant reasons. First, we recognize that the indigenous peoples of the Americas are affected by arthritis at a disproportionate rate than other racial and ethnic groups. Secondly, we acknowledge that there is a history of erasure of the advances made by indigenous scientists and knowledge seekers. Third, because we are taught that by speaking the language of our ancestors that we breathe life into it and strengthen all our relations. We finally acknowledge that the research in this study was primarily performed on the traditional territories and ancestral homelands of the Cheyenne, Arapaho, and Ute nations. This area, specifically the confluence of the Platte and Cherry Creek Rivers was the epicenter for trade, information sharing, planning for the future, community, family and ally building, as well as conducting healing ceremonies for over 45 Indigenous Nations, including the Lakota, Kiowa, Comanche, Apache, Shoshone, Paiute, Zuni, Hopi among others.

## List of Supplementary Materials

### Supplemental Methods

Supplemental Table 3: <u>Plasmablast Cohort RA-relevant antigen binding full</u>

Supplemental Table 7: <u>Variable region sequences for expressed plasmablast antibodies</u>

## Detailed Experimental Methods

### Human Subjects

Participants included in this study were selected from the Studies of the Etiology of Rheumatoid Arthritis (SERA) cohort, described previously(*78*) and in detail in Supplemental Methods. Briefly, SERA is a cross-sectional and longitudinal study that follows and studies individuals at-risk for future development of RA. At-risk status is defined as individuals with positive levels of circulating RA-specific autoantibodies (serum anti-CCP3/anti-CCP3.1 positive and/or ≥2 RF isotypes), or first-degree relatives (FDRs) of a patient with RA. All at-risk individuals were without a history or findings of inflammatory arthritis at the time of sample acquisition. Early RA (eRA) was defined as having received a diagnosis of seropositive RA within the past 12 months. Controls were defined as having a negative CCP3, CCP3.1, and RF; not having a FDR with RA or a personal history of autoimmune rheumatic disease; and no inflammatory arthritis by history or on exam at the time of the study.

Participants were recruited at the University of Colorado Anschutz Medical Campus with approval from the Colorado Multiple Institutional Review Board (IRB# 01-675). During study visits, all participants were asked to complete questionnaires that assessed basic demographics, self and family history of disease, and past and current environmental exposures. A 68-joint examination was performed by a rheumatologist or trained study nurse in the at-risk and control subjects to confirm the absence of inflammatory arthritis. Blood was drawn into serum separation tubes (Fisher Scientific BD Vacutainer^TM^), allowed to clot for 15 minutes at room temperature and then centrifuged at 3000 rpm for 10 minutes. A fecal collection kit (BD GasPak EZ Anaerobe Pouch System with Indicator) was provided to the participant for self-collection and overnight shipment on frozen gel packs via FedEx to the study site. Serum and fecal samples were aliquoted into 2 ml graduated tubes and stored at −80°C until the sample was utilized for further analysis.

### Serum Autoantibody Testing

Anti-CCP3 (IgG, Invoa Diagnostics) and anti-CCP3.1 (IgG/IgA, Inova Diagnostics) ELISAs were performed and analyzed according to the manufacturers’ instructions with results reported in units/mL. RF IgG and IgA isotypes were measured by ELISA using QUANTA Lite kits (Inova Diagnostics), and results are reported in international units/mL.

### Human Plasmablast Monoclonal Antibody Expression

As described previously(*32*), dual IgA/IgG family plasmablasts (n=94) were isolated from 4 individuals at-risk for RA, defined as serum RF+ only (n=1), anti-CCP+ only (n=1) and RF and anti-CCP+ (n=2), as well as from 2 anti-CCP+ individuals with early RA. Determination of plasmablasts belonging to a clonal family was based on shared, IMGT-based assignments of HC and LC V-J-region genes and 60% identity matching. A fraction of these plasmablasts were selected for cloning and immortalization due to their representation of shared IgG/IgA clonal families or their ability to bind citrullinated peptide tetramers.

Production of the monoclonal antibodies was performed using an Expi293 Expression System (ThermoFisher) with Expi293F cells, as previously described(*79–81*). Fab domains from the plasmablast-derived mAbs (PB-mAbs), selected for binding of RA-relevant antigens on a protein microarray, were expressed in a mouse IgG2a scaffold. Antibodies were recombinantly produced in Dr. Robinson’s laboratory at Stanford University or by staff at LakePharma. The variable region sequences of these antibodies are shown (Supp. Table 7).

### RA-Related Autoantigen Array

Titers of IgG antibodies targeting RA-associated autoantigens (n=346) were measured using a previously described Bio-Plex assay(*82*). Analysis was performed on a Luminex 200 instrument running Bio-Plex Manager software version 6.1. 94 plasmablast-associated mAbs isolated from six human subjects (n=4 at-risk for RA and n=2 early RA) were analyzed. Subsequently thirty mice were analyzed across two timepoints utilizing this approach.

### Fecal Bacteria Pool

Feces from 5 controls, 8 at-risk (n=3 non-FDR CCP+, n=2 FDR CCP+, n=3 FDR CCP-), and 5 eRA individuals were utilized to create a pool. Inclusion of samples was based on fecal 16S sequencing data from these individuals, with a goal of being broadly representative of bacteria across all study groups. 50 mg of fecal material from each sample was placed onto ceramic bead tubes (MP Biomedicals) and were rehydrated in 1mL sterile PBS. Each sample was homogenized by bead beating for 15 minutes at full speed. Samples were then centrifuged at 50 x g to remove debris, and the supernatants from each sample was collected into a combined tube. The combined bacteria were washed 3 times with sterile PBS containing 1% (w/v) BSA (Sigma-Aldrich) and spun at 14,000 x g to pellet bacteria. In order to verify the bacterial composition of this pool, DNA was extracted and 16S rRNA sequencing was performed per the methods below.

### Plasmablast-Bound Bacteria Sequencing

100 uL of fecal bacteria pool (400 mg feces/mL), or 5 x 10^6^ CFU cultured bacteria, were combined with 0.5 ug of PB-mAbs, and was incubated on ice for 30 min. The samples were washed 3 times with sterile PBS containing 1% (w/v) BSA. 0.5 ug of secondary antibody (PE Rat anti-mouse IgG2a, Invitrogen) was added to each sample, along with 1:4000 nucleic acid dye Syto9Green (Invitrogen). Negative controls included in-house generated PB-mAbs against the 2010/2011 seasonal trivalent influenza vaccine (H1N1 A/California/7/2009, H3N2 A/Perth/16/2009, and B/Brisbane/60/2008)(*83*) and *Borrelia burgdorferi.* The positive control was polyclonal goat anti-*E. coli* antibody (Invitrogen)with PE mouse anti-goat (SouthernBiotech) secondary. Each sample was incubated again for 30 minutes on ice and underwent the same washing step. Samples were analyzed by flow cytometry on a BD LSR2 and FlowJo v10 software. A positive mAb-bacteria binding cut-off was established as >2 SD above the mean fluorescence intensity of the negative controls. Those mAbs with positive binding then underwent subsequent flow cytometric sorting into mAb coated and uncoated fractions utilizing an Astrios EQ 5 laser flow sorter (Beckman Coulter). Sorted bacterial DNA was extracted and 16S rRNA sequenced.

### DNA Extraction

50 mg of fecal material from each sample was placed onto ceramic bead tubes (MP Biomedicals) with 1mL sterile PBS. For samples that had undergone flow sorting, the mAb-coated and uncoated fractions were spun at 14,000 x g for 30 minutes, and supernatant was removed to create a total sample volume of 100 uL that was added to ceramic bead tubes. Samples were homogenized by bead beating for 10 minutes at maximum speed and then centrifuged at 50 x g for 15 min at 4°C to remove debris. Supernatants were harvested and centrifuged at 14 000 x g for 15 min at 4°C to pellet the bacteria. Supernatant was removed, and the bacteria were washed 3 times with sterile PBS containing 1% (w/v) BSA (Sigma-Aldrich) and spun at 14 000 x g to pellet bacteria. Total bacterial genomic DNA extractions were performed using the Qiamp Power Fecal DNA prep kit (Qiagen) according to the manufacturer’s instructions.

### 16S Sequencing and Analysis

DNA was amplified by polymerase chain reaction (PCR) with broad-range bacterial primers targeting the 16S ribosomal RNA (rRNA) gene hypervariable regions V1 and V2 and pooled amplicons were subjected to Illumina MiSeq sequencing, as previously described(*84–86*). Assembled sequences were aligned and classified with SINA (1.3.0-r23838)(*87*) using the 418,497 bacterial sequences in Silva 115NR99(*88*) as reference configured to yield the Silva taxonomy. Operational taxonomic units (OTUs) were produced by clustering sequences with identical taxonomic assignments. OTUs with >0.01% abundance in any sample and observed in >5% of the samples were included in further analyses. Analyses of OTU relative abundance and biodiversity were conducted using Explicet software(*89*). Microbiome data were analyzed using Explicet and R statistical software, including MicrobiomeAnalyst(*90*). Tests of overall community composition used a non-parametric multiple analysis of variance (PERMANOVA) test of Morisita-Horn dissimilatiry scores, with p-values estimated through 1,000,000 permutations. Alpha-diversity indices were estimated through 1,000 resamplings at the rarefaction point of 90,000 sequences. Between-group differences in alpha-diversity were assessed by ANOVA.

### Bacterial Isolation

Primary bacterial isolates were established from a fecal sample of an at-risk individual by serially diluting the stool sample 1:10 in sterile oxygen-reduced Mega media(*91*), adding 1 g of feces to 10mL of media, in a Coy anaerobic chamber. The diluted sample was homogenized for 5 minutes by vortexing, and then solids were allowed to settle for 5 minutes. Serial 1:10 dilutions were carried out to 10^-9^ dilutions in Mega media. For the 10^-7^, 10^-8^, and 10^-9^ dilutions, 170 uL of media was plated in each well of a 96 well plate. Each 96 well plate was sealed and allowed to incubate in anaerobic conditions at 37°C for 5 days. The dilution factors that resulted in 30% or fewer wells being turbid at the end of the incubation had less than a 5% chance of having multiple bacterial clones in each well. To ensure strain purity, each established isolate was cultured on a sheep blood agar (SBA) plate under anaerobic conditions for 3 days and then had one isolated colony harvested and grown to larger volume in sterile Mega media under anaerobic conditions. Each isolate was stored at -80°C in 25% sterile oxygen-reduced glycerol.

### Taxonomic Identification

Each bacterial isolate was taxonomically identified using bacterial group primers for *Lachnospiraceae* and *Ruminococcaceae*. The primer sequences were 5’è3’ Forward: CGGTACCTGACTAAGAAGC and Reverse: AGTTT(C/T)ATTCTTGCGAACG as compared to global 16S bacterial primers, which were 5’ è 3’ rpoB1698 Forward: AACATCGGTTTGATCAAC and rpoB2041 Reverse: CGTTGCATGTTGGTACCCAT. The cycling protocol is as follows: 50°C for 2 minutes, 95°C for 10 minutes, and then 40 cycles of 95°C for 15 seconds to 60°C for 1 minute. Isolates whose DNA amplified with the *Lachnospiraceae/Ruminococcaceae* primers were further 16S sequenced for confirmation. Confirmed isolates were subsequently whole genome sequenced by Novogene, Inc. using a 350 bp insert DNA library and an Illumina Platform PE150. Short reads were cleaned and contigs and scaffolds assembled using Abyss(*33*). Long reads were verified as derived from order *Clostridiales* through NCBI-BLAST. Assembled long reads were aligned to a reference genome, *Clostridiales* strain MGYG-HGUT-02424 (Genome accession number GCF_902387115.1, located at https://ftp.ncbi.nlm.nih.gov/genomes/all/GCF/902/387/115/GCF_902387115.1_UHGG_MGYG-HGUT-02424/) and analyzed using Artemis(*92*).

### Mice

DBA/1j mice were obtained from The Jackson Laboratory and derived germ-free by Taconic via embryonic transfer into germ-free Swiss Webster hosts. Pregnant dams were shipped to the University of Colorado Anschutz Medical Campus Gnotobiotic Core facility, where a colony of germ-free DBA/1 mice was maintained in sterile vinyl isolators. Mice were age and sex matched for all studies. Between 6-10 weeks of age (mean=8.2), mice were orally gavaged with 10^7^ CFU in 200 μl of primary *Subdoligranulum* species isolates 1 or 7, *Prevotella copri* (DSM 18205), or sterile growth media. Mono-colonized mice were monitored weekly for stable bacterial colonization; fecal pellets were collected, DNA was extracted, and each sample was amplified using global 16S primers. Furthermore, cecal contents collected at termination were serially diluted and plated on SBA and incubated for 48 hours at 37°C in anaerobic conditions. Bacterial concentrations were determined by quantifying the number of CFUs at each dilution factor.

Mice were monitored for arthritis development similar to that performed for the collagen-induced arthritis (CIA) model(*10*) . Arthritis severity was measured as a mean clinical score for each of the animal’s four paws, where 0 = normal, 1 = erythema, 2 = swelling, and 3 = ankylosis. At the specified time points, feces, serum, and tissues were harvested from mice. All animal experiments were approved by the University of Colorado School of Medicine Institutional Animal Care and Use Committee.

### FITC-Dextran Flux

Mice were orally gavaged with 0.6 mg/kg body weight 4 kDa dextran labeled with fluorescein (Sigma) and serum was collected 4 hours later. The amount of fluorescence was measured with a fluorimeter (Promega) at 485/530 nm. A standard curve was generated to calculate the amount of dextran that was present in the sera.

### Histopathology

Whole colon and ileal tissue was harvested from mice, flushed with PBS, dissected longitudinally, and pinned in wax for fixation in formaldehyde overnight. Tissues were then embedded in paraffin. Sections of 5 μm were cut and stained with hematoxylin and eosin (H & E). Four high-powered fields of well-oriented colon tissue per mouse were analyzed at 40X magnification for quantification of tertiary lymphoid structures, crypts per high-powered field, and crypt height and width.

Mouse paws were removed at mid-limb and fixed in 10% paraformaldehyde. The bones were decalcified in 14% EDTA for 10 weeks and then embedded in paraffin. Sections of 5um were cut from paraffin embedded tissues and stained with hematoxylin and eosin (H & E). Pathology was assessed by a trained pathologist for synovitis, osteomyelitis, vasculitis, subcutaneous/muscle/periosteal inflammation, and fat inflammation, and global incidence rate of pathological findings were reported.

### Complement C3 Immunohistochemistry

Paraffin-embedded tissue slides were assessed for C3 complement deposition in the joints. Each slide was exposed to the following: xylene (histology grade, Sigma) for 5 minutes (x2), 100% ethanol for 1 minute (x2), 95% ethanol for 1 minute (x2), tap water for 2 minutes, wash buffer (Dako Cytomation) for 2 minutes, 3% hydrogen peroxide (Millipore-Sigma) for 5 minutes, wash buffer for 2 minutes (x2), serum-free protein block (Dako Cytomation) for 5 minutes, goat anti-mouse complement C3 (1:10,000) (Fisher) overnight at 4°C, wash buffer for 2 minutes (x2), goat probe (BioCare Medical) for 15 minutes, wash Buffer for 2 minutes (x2), goat polymer (BioCare Medical) for 15 minutes, wash buffer for 2 minutes (x2), DAB+ (Dako Cytomation) for 5 minutes, distilled water for 2 minutes (x2), hematoxylin (4-5 dips), distilled water for 1 minute, 95% ethanol for 1 minute, 100% ethanol for 1 minute (x2), xylene for 1 minute (x2), and then were mounted and coverslipped. Each stained section was assessed for complement deposition in the joint and was scored in a blinded fashion from 0-3 based on C3 deposition severity: 0= no staining, 1= mild staining, 2= moderate staining, 3=intense staining.

### IgG and IgA Immunohistochemistry

Paraffin-embedded tissue slides were assessed for IgA/IgG deposition in the joints. Each slide was exposed to the following: xylene (histology grade, Sigma) for 5 minutes (x2), 100% ethanol for 1 minute (x2), 95% ethanol for 1 minute (x2), tap water for 2 minutes, wash buffer (Dako Cytomation) for 2 minutes, 3% hydrogen peroxide (Millipore-Sigma) for 5 minutes, wash buffer for 2 minutes (x2), serum-free protein block (Dako Cytomation) for 5 minutes, anti-mouse IgA-HRP 1:500 or anti-mouse IgG-HRP 1:500, respectively, wash buffer for 2 minutes (x2), DAB+ (Dako Cytomation) for 5 minutes, distilled water for 2 minutes (x2), hematoxylin (4-5 dips), distilled water for 1 minute, 95% ethanol for 1 minute, 100% ethanol for 1 minute (x2), xylene for 1 minute (x2), and then were mounted and coverslipped. Each stained section was assessed for IgA/IgG deposition in the joint and was scored in a blinded fashion from 0-3 based on IgA/IgG deposition severity. 0= no staining, 1= mild staining, 2= moderate staining, 3=intense staining.

### Immunophenotyping Flow Cytometry

Spleen, mesenteric lymph nodes (MLNs), and Peyer’s patches (PPs) were harvested from monocolonized mice at days 14 and 35 following gavage. Tissues were processed by homogenizing in RPMI media and straining through a 70 micron filter. Cells were pelleted by centrifugation at 500xg for 8 minutes. Splenic cells underwent RBC lysis using a commercially available buffer (eBioscience) for 10 minutes and were then centrifuged again. Cells were then resuspended in 1mL FACS buffer (PBS with 5% FBS) for cell counting. 10^6^ cells were added to each staining tube. For intracellular stains, cells were permeabilized and fixed using a commercially available buffer (Tonbo). Supplemental Table 1 lists the antibodies and clones used (Supplemental Table 8). The panel was validated through antibody titration, full minus one (FMO) controls, and isotype controls. Cytometric analysis was performed on a Cytek spectral cytometer, and downstream analysis was performed using FlowJo v10 software.

### Serum Transfer and Analysis of Arthritis

Serum collected from mono-colonized and control mice was pooled from 10 mice for each treatment group, respectively, and 150 μl injected IP into germ-free DBA/1 mice, as we know the background autoantibody levels for these mice. The mice were then monitored daily for the development of arthritis and scored in a blinded fashion as above. To determine the kinetics of the joint swelling, mice were monitored for 21 days post-serum transfer. The peak of joint swelling was determined to be 7 days post-serum transfer. Then, a second cohort of mice were dosed with serum, and were sacrificed 7 days following serum transfer and paws were collected for histology.

### Lymphocyte and Granulocyte Depletion Studies

CD4+ T cells, B cells, and neutrophils were depleted from germ-free DBA/1 mice by injection of targeting monoclonal antibodies. For CD4+ T cell depletion, mice were injected intra-peritoneally with 200 μg of antibody GK1.5 (BioXCell) two days prior to colonization, and then with 100 μg of antibody IP every four days thereafter. Mice were depleted of B cells using a CD20 depletion antibody (clone SA271G2, Biolegend). Mice were injected with 250 μg of antibody IP two days before bacterial administration, and then with 250 μg of antibody IP on day 21 of the study. Neutrophils and other myeloid-derived cells were depleted using a Ly6G/Ly6C antibody (clone GR1, BioXCell) injected IP with 200 μg two days before bacterial administration, and then with 200 μg of antibody IP every four days thereafter. Finally, 200 μg of control antibody (anti-Rat IgG2aK, clone RTK4530, Biolegend) was injected IP two days before bacterial administration, and then with 200 μg of antibody IP on day 21 of the study. Cellular depletion was confirmed in the MLNs, PPs, and spleen of mice. The depletions were efficacious at depleting the B cells by 2-fold, the T cells by 10-fold, and granulocytes by 5-fold on average. Mice were gavaged with 10^7^ CFU of *Ruminococcaceae subdoligranulum* isolate 7 on the second day of the study and monitored for the development of arthritis for 35 days. Mice were then sacrificed and tissues harvested.

### Human T cell studies

Healthy and RA participants were recruited with informed consent through the Benaroya Research Institute (BRI) rheumatic disease registry. Sample use was approved and monitored by the BRI Institutional Review Board (IRB number: IRB07109-139). All participants met the 1987 American College of Rheumatology criteria. Clinical characteristics of the participants are summarized in Supplemental Tables 5 and 6.

PBMC were isolated from whole blood by the BRI Clinical Core Laboratory. Briefly, PBMC were separated from whole blood over Ficoll-Hypaque gradient, cryopreserved in heat inactivated FBS supplemented with 10% DMSO, and stored in liquid nitrogen. PBMC were thawed in a 37°C water bath, washed in RPMI media supplemented with 10% FBS and 0.001% DNase/Benzonase (Sigma Aldrich), re-suspended in serum-free RPMI at a volume of 10^7^ cells/mL, and allowed to rest in 37°C 5%-CO_2_ incubator for 2 hours. Cells then were centrifuged, re-suspended in RPMI supplemented with 10% commercial human pooled serum and anti-CD40 antibody (Miltenyi) at 1µl/million cells, and plated at a volume of 500µl of 5 million PBMC per well of a 48-well tissue culture plate. PBMC were stimulated 14 hours in 37°C in the presence of either 0.1% DMSO or 50ng/mL isolate 7 or isolate 1, respectively. For class II blocking, PBMC were rested in either anti-HLADR (clone L243) at 20ug/mL, or equal volume PBS, 30 minutes prior to stimulation.

After 14-hour stimulation, cells were transferred to a 4 mL polypropylene FACS tube for surface stain with PE-CF594 mouse anti-CD154 (PECF594, BD) and PE-Cy7 mouse anti-CD137 (CY1G4, Biolegend)for 15 minutes RT, then enriched with anti-PE microbeads (Miltenyi) following manufacturer’s standard protocol with 1% of PBMC set aside before enrichment to calculate number of CD4 cells present. Cells were then surface stained for 30 minutes at 4°C with the following antibodies: CD14 FITC, CD19 FITC, CD56 FITC, CD3 PerCPCy5.5, CD4 BV510, CD8 APCCy7, CD69 AF647, CD45RA BV421, and CCR7 BV711 (Supp. Table 9). Cells were then stained with Sytox green (Thermo) to exclude non-viable cells prior to FACS analysis on a BD FACS Canto II. Flow cytometry data were analyzed using FlowJo v10, SAS JMP statistical software V15, and Graphpad Prism 8.0. The frequency of activated CD4 T cells was calculated as follows: F = (1,000,000 x activated events) / (100 x number of CD4 T cell events from the pre-enriched fraction).

---

**Supplemental Figure 1:**
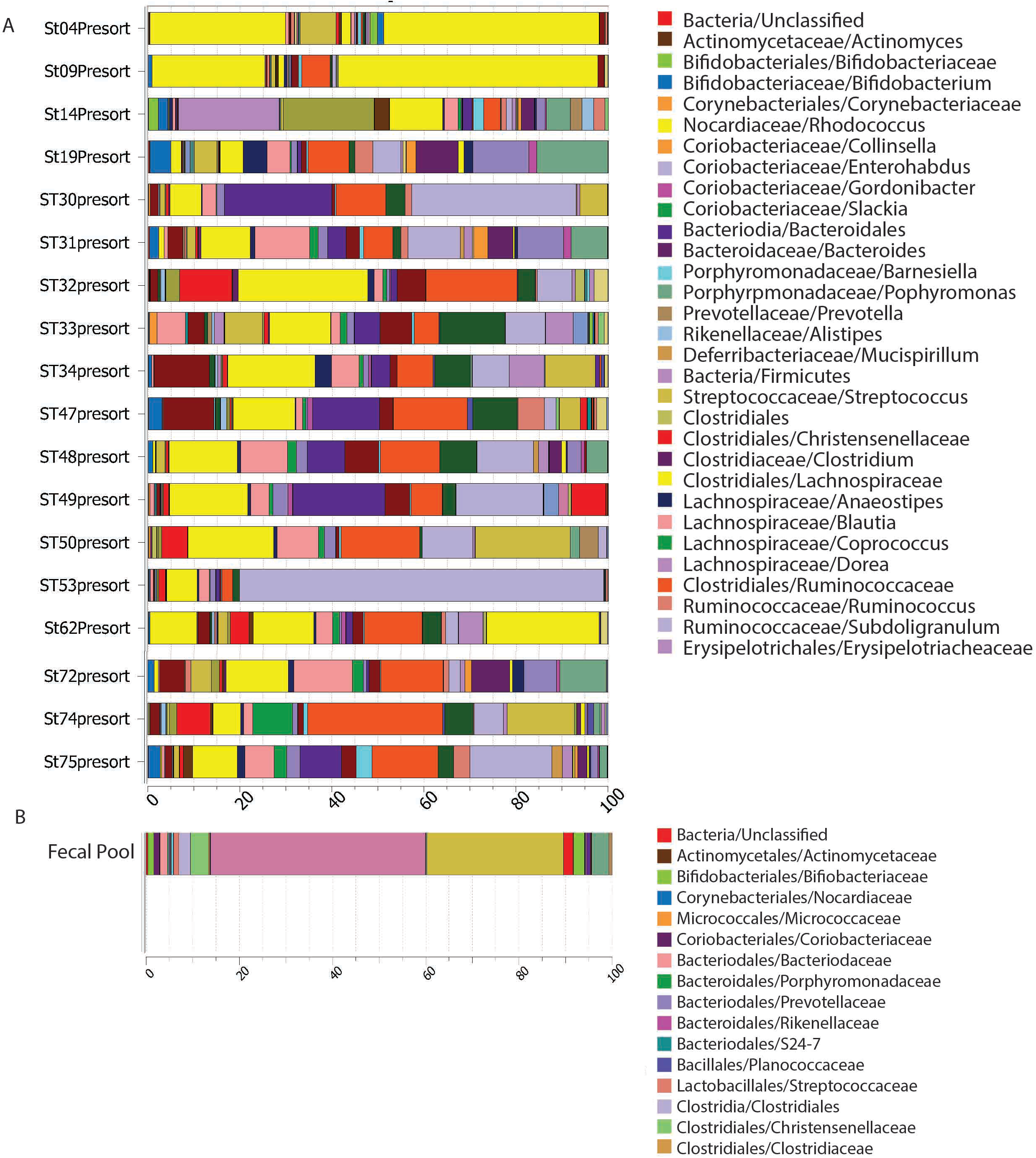
l6S sequencing of each fecal pool sample. Fecal samples were collected from healthy control individuals (n=5; ST30, 31, 32, 33, and 34), individuals at-risk for RA (n=8; ST4, 9, 14, 19, 62, 72, 74, and 75), and individuals with early RA (n=5; ST 47, 48, 49, 50, and 53, <l year from diagnosis). Each sample was individually 16S sequenced and the OTU table graphed in a stacked bar format **(A).** Tue most highly represented taxonomic groups are represented (see figure legend to the right), displaying the percentage of the bacteria sequenced belonging to each taxa (x-axis) for each sample (y-axis). A pool of fecal bacteria was created from these human fecal samples utilizing The bacteria pool was 16S sequenced and its bacterial components are displayed in a bar format. The most highly represented taxonomic groups are represented (see figure legend to the right), displaying the percentage of the bacteria sequenced belonging to each taxa (x-axis) for the combined fecal bacterial pool **(B).**

**Supplemental Figure 2:**
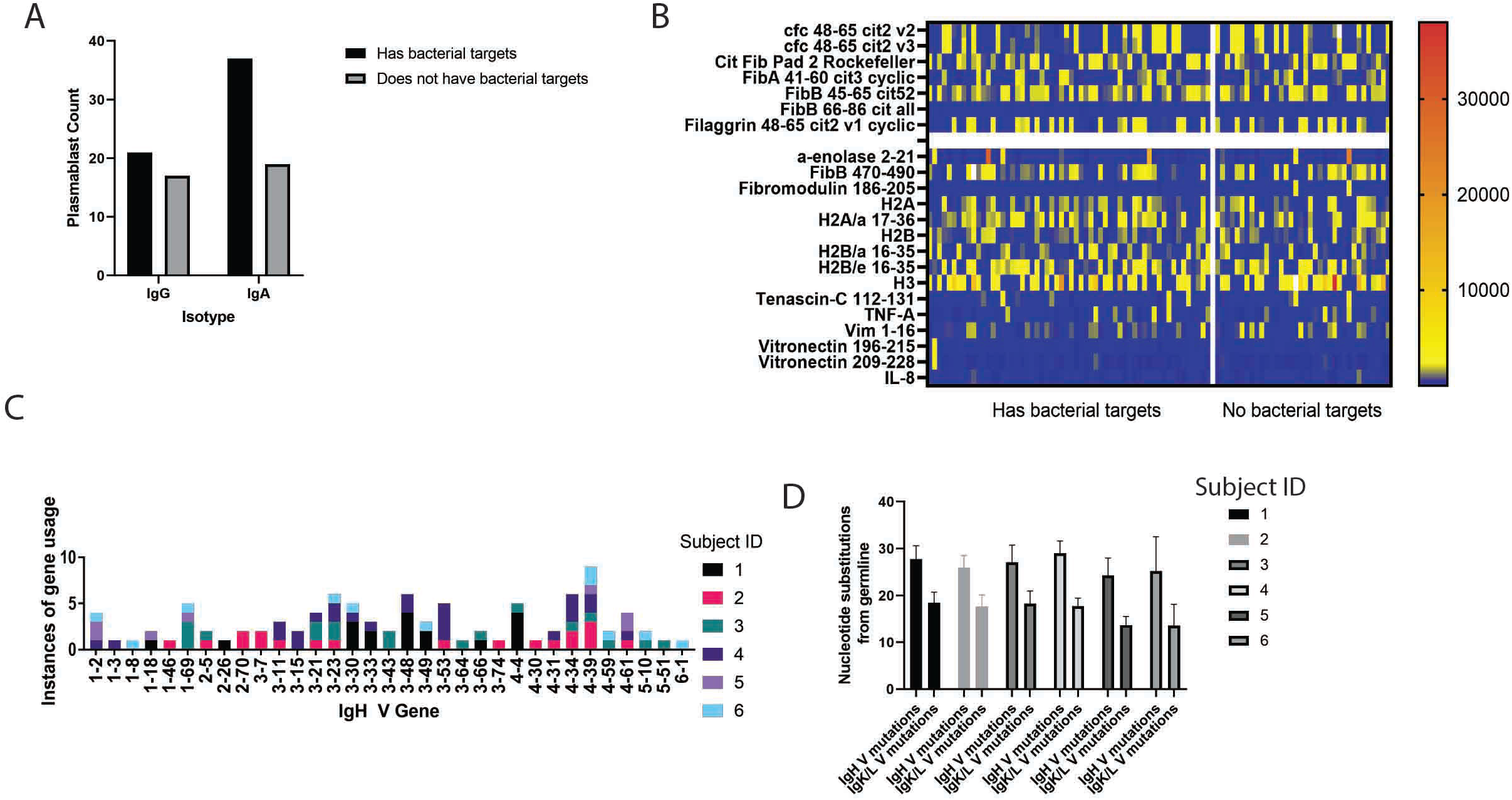
Plasmablast cohort Vh gene usage, mutations from germline, and bacterial binding characteristics. **(A)** Displayed is the isotype of the each plasmablast antibody (lgG or IgA, x-axis) segregated by whether or not they target intestinal bacteria, with mAbs targeting bacteria shown in black bars and mAbs without bacterial targets shown in grey bars. The total mAb count for each group is displayed on the y-axis. Fisher’s exact test p=0.38 **(B)** RA-relevant autoantigenic targets are displayed as in Fig. IA but segregated by whether the plasmablasts (listed along the X-axis) have bacterial binding targets or not. 94 plasmablast-derived mAbs from at-risk (n=4) and eRA (n=2) individuals belonging to shared IgG and IgA clonal families were applied to a planar array containing 346 different citrullinated and native peptide targets. The heatmap demonstrates degree of reactivity between PB-mAbs that have bacterial targets (x­axis, left) and PB-mAbs that don’t have bacterial targets (x-axis, right) with specific antigens (y-axis). **(C)** The IgH V gene used by each plasmablast is displayed by individual from whom the PB-mAb was derived. The instances of gene usage is displayed on the x-axis, with the total mAb count displayed on the y-axis. **(D)** Instances of nucleotide substitutions from gerrnline (y-axis) are demonstrated for each participant, as shown in the figure legend (x-axis). IgH V mutations are displayed on the left and IgL/K mutations are displayed on the right for each subject.

**Supplemental Figure 3:**
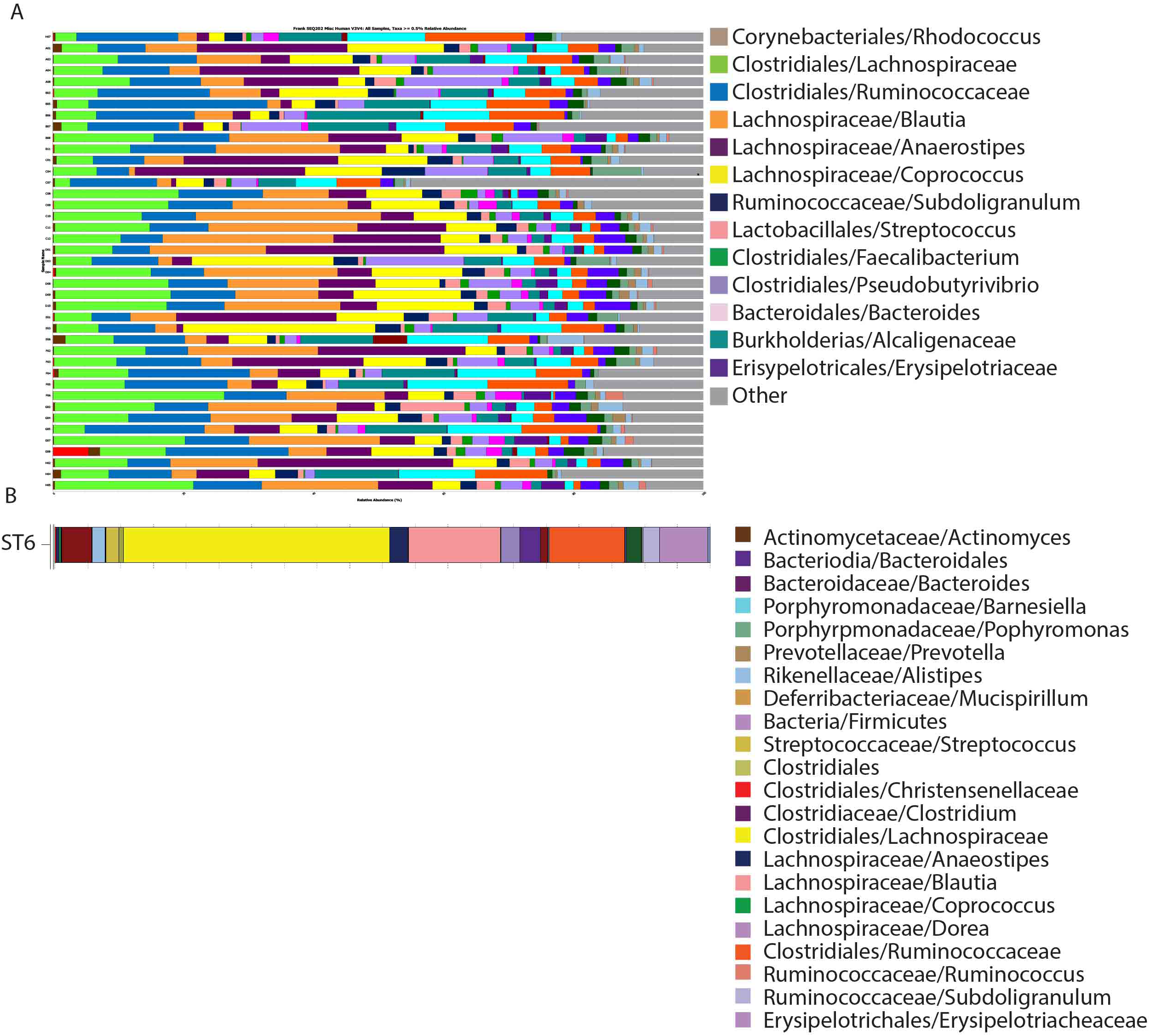
Plasmablast mAbs bind bacteria from a broadly representative fecal pool. **(A)** The ability of the plasmablasts to bind to the bacteria in the fecal pool was detennined by flow cytometry. Each mAb was bound to the fecal pool; the bound bacterial fraction was separated by flow cytometric sorting and each bound fraction was 16S rRNA sequenced. Each sample was individually 16S sequenced and the OTU table graphed in a stacked bar format. The most highly represented taxonomic groups are represented (see figure legend to the right), displaying the percentage of the bacteria sequenced belonging to each taxa (x-axis) for each discrete mAb that bound bacteria (y-axis). **(B)** The sample from which bacterial strains were isolated was 16S rRNA sequences and the OTU table graphed in a stacked bar format.

**Supplemental Figure 4:**
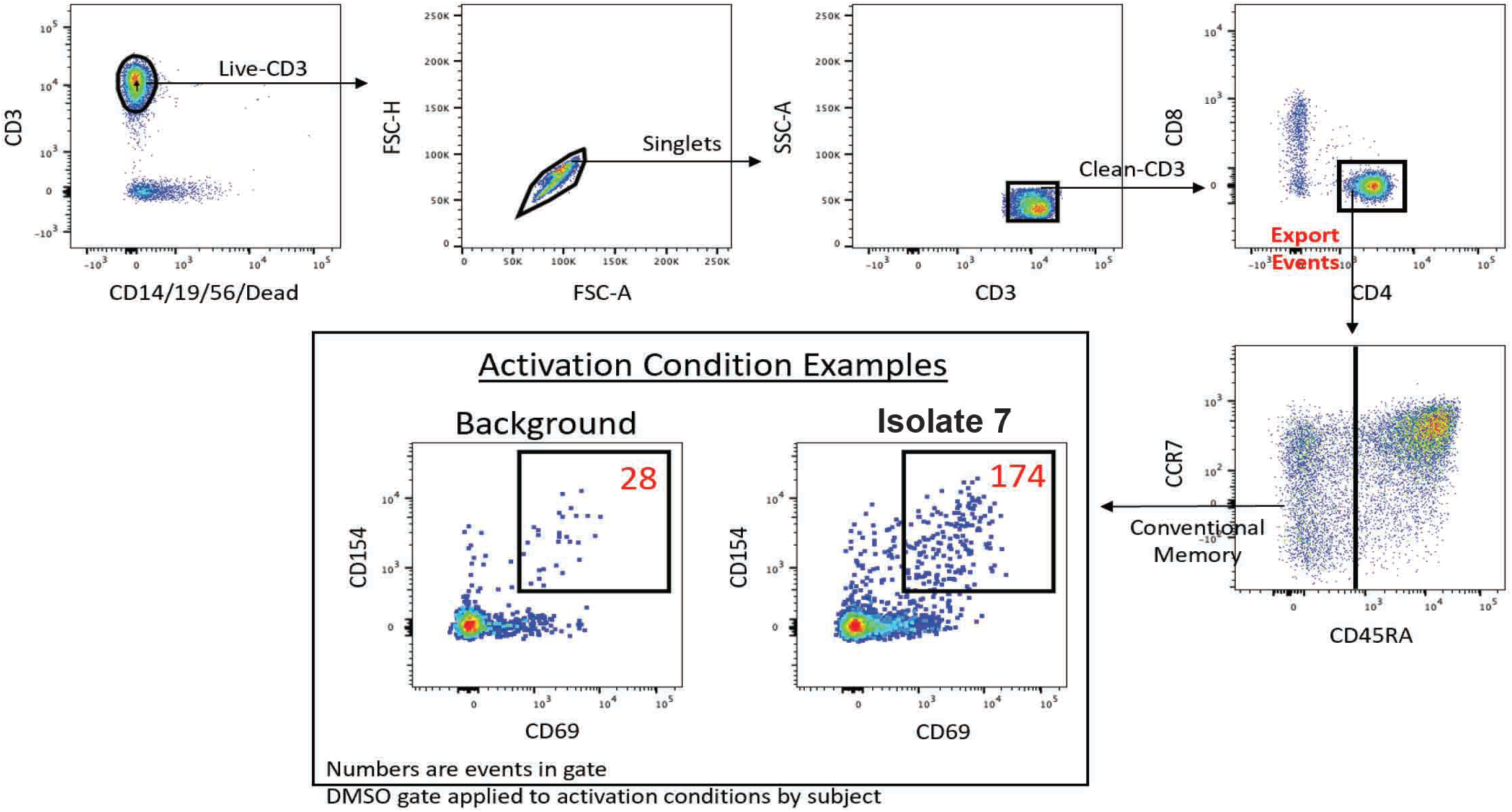
Gating strategy and validation of stimulated CD4+ T cells. PBMCs from RA cases (n=11) were surface stained prior to stimulation for 30 minutes at 4°C with the following antibodies: CD14 (Biolegend)/CD19 (Biolegend)/CD56 (BD) FITC, CD3 PerCPCy5.5, CD4 BV510, CD8 APCCy7, CD69 AF647, CD45RA BV421, and CCR7 BV711 (all Biolegend). Cells were then stained with Sytox green to exclude non-viable cells prior to FACS analysis on a BD FACS Canto II. The gating scheme for this flow cytometric panel is shown, with arrows representing the direction of subsequent gating events.\

**Supplemental Figure 5:**
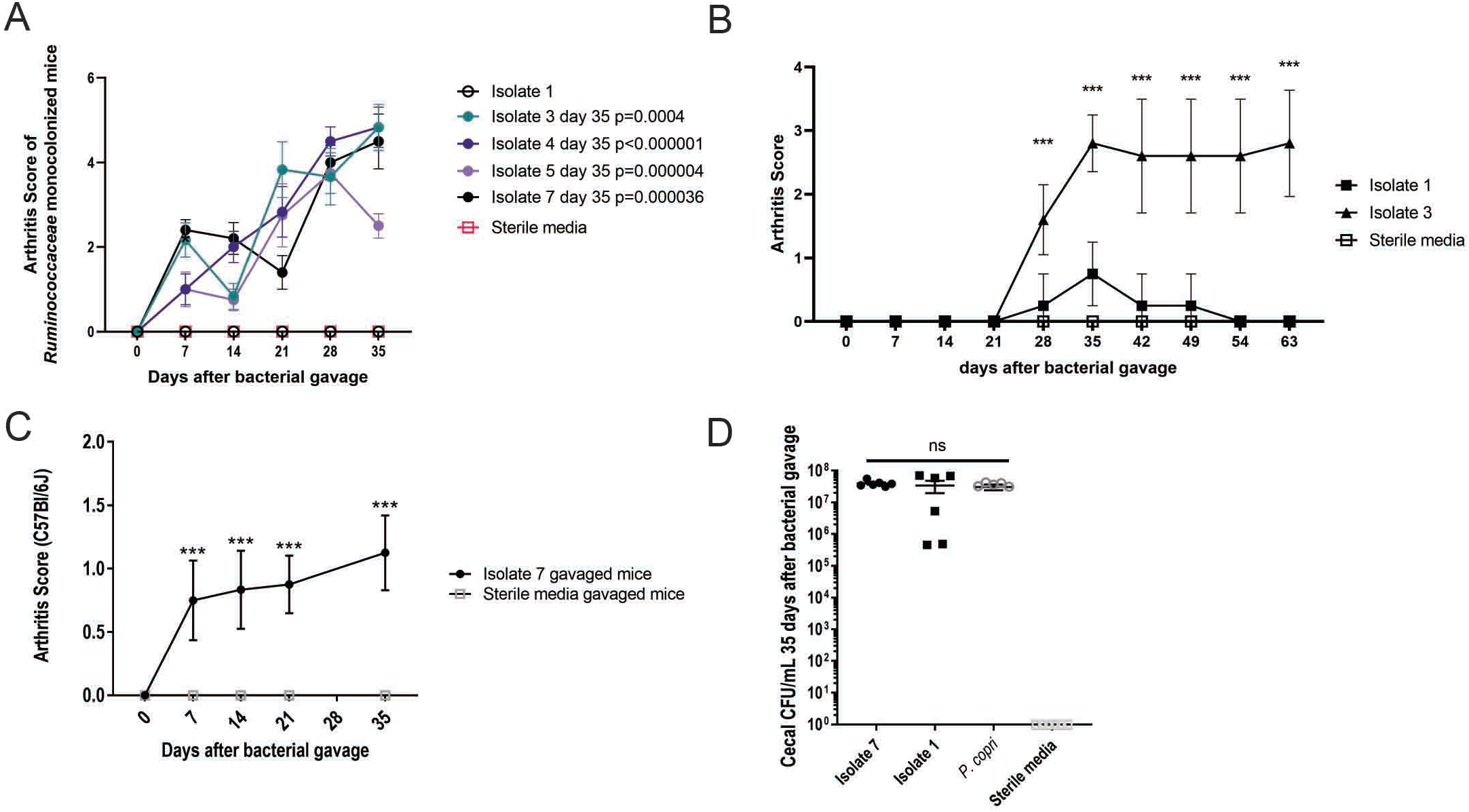
Mono-colonization of germ free mice with *Ruminococcaceae* isolates results in a spectrum of joint swelling phenotypes. **(A)** *Ruminococcaceae Subdoligranulum* Isolates 1, 3, 4, 5, & 7 were gavaged separately into germ-free DBA/1 mice. Mice were gavaged with either 5 x 10^6^ CFUs of the cultured bacteria or sterile media. N=6 mice gavaged with isolate 1, n=6 with isolate 3, n=6 with isolate 4, n=4 with isolate 5, n=5 with isolate 7, and n=6 with sterile media. The mice were then observed weekly for 35 days for the development of joint swelling or ankylosis. The clinical score is shown (y-axis) relative to time post-bacterial gavage (x-axis) for each different isolate. (p-values for each of the isolates relative to sterile media gavaged control mice at day 35 is shown in the figure legend, unpaired t-test). **(B).** *Ruminococcaceae Subdoligranulum* Isolates 1 & 3 were gavaged separately into germ-free DBA/1 mice. Mice were gavaged with either 5 x 10^6^ CFUs of the cultured bacteria or sterile media (n=4 isolate l gavaged, n=5 isolate 3 gavaged, and n=5 sterile media gavaged mice). The mice were then observed weekly for 63 days for the development and maintenance of joint swelling or ankylosis. The clinical score is shown (y-axis) relative to time (x-axis) for each isolate.***, P<0.001, multiple unpaired t-test. **(C)** *Ruminococcaceae Subdoligranulum* isolate 7 was gavaged into germ-free C57Bl/6J mice. Mice were gavaged with either 5 x 10^6^ CFUs of the cultured bacteria (n=8) or sterile media (n=8). The mice were then observed weekly for 35 days for the development of joint swelling or ankylosis. The clinical score is shown (y-axis) relative to time (x=axis) for each treatment group.***, P<0.001 at day 35, unpaired t-test. **(D)** To verify stable colonization of the monocolonized mice, the ceca of mice were harvested and CFU/mL of bacteria was determined (n=6 mice from each group, respectively). The total CFU/mL at day 35 is displayed (y-axis) relative to each treatment group (x-axis). P>0.95 for isolate groups compared to each other, one-way ANOVA.

**Supplemental Figure 6:**
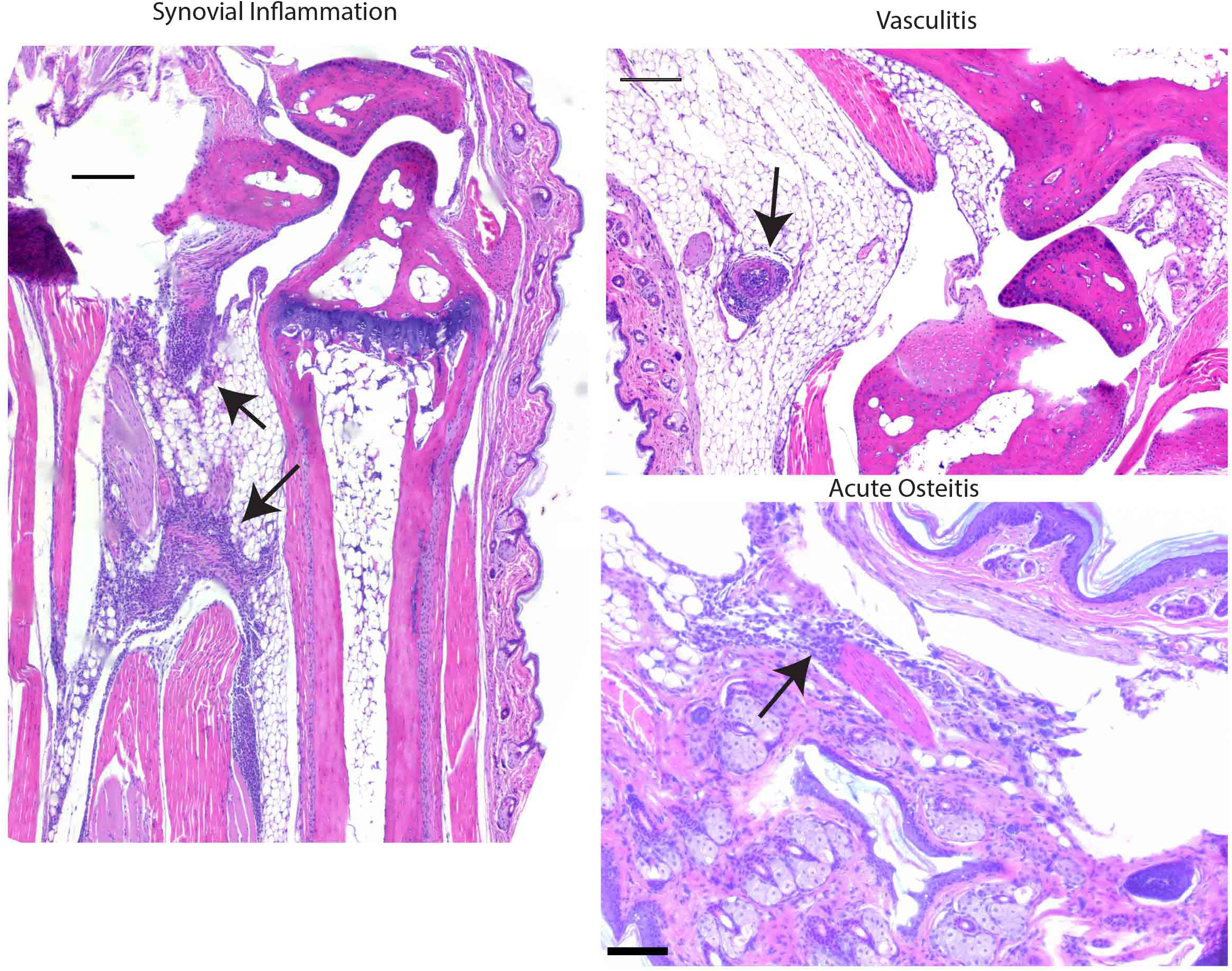
Mono-colonized mouse paw pathology. H&E staining was performed on decalcified paw sections from isolate 7, isolate 1, *P. copri,* and sterile media gavaged mice and paw pathology was assessed by a pathologist in a blinded fashion. Evaluation of pathology included synovitis (top left photo, arrows), vasculitis (top right photo, arrow), and acute osteitis (bottom right photo, arrow) **in** isolate 7 monocolonized mice. Scale bar is lOOum.

**Supplemental Figure 7:**
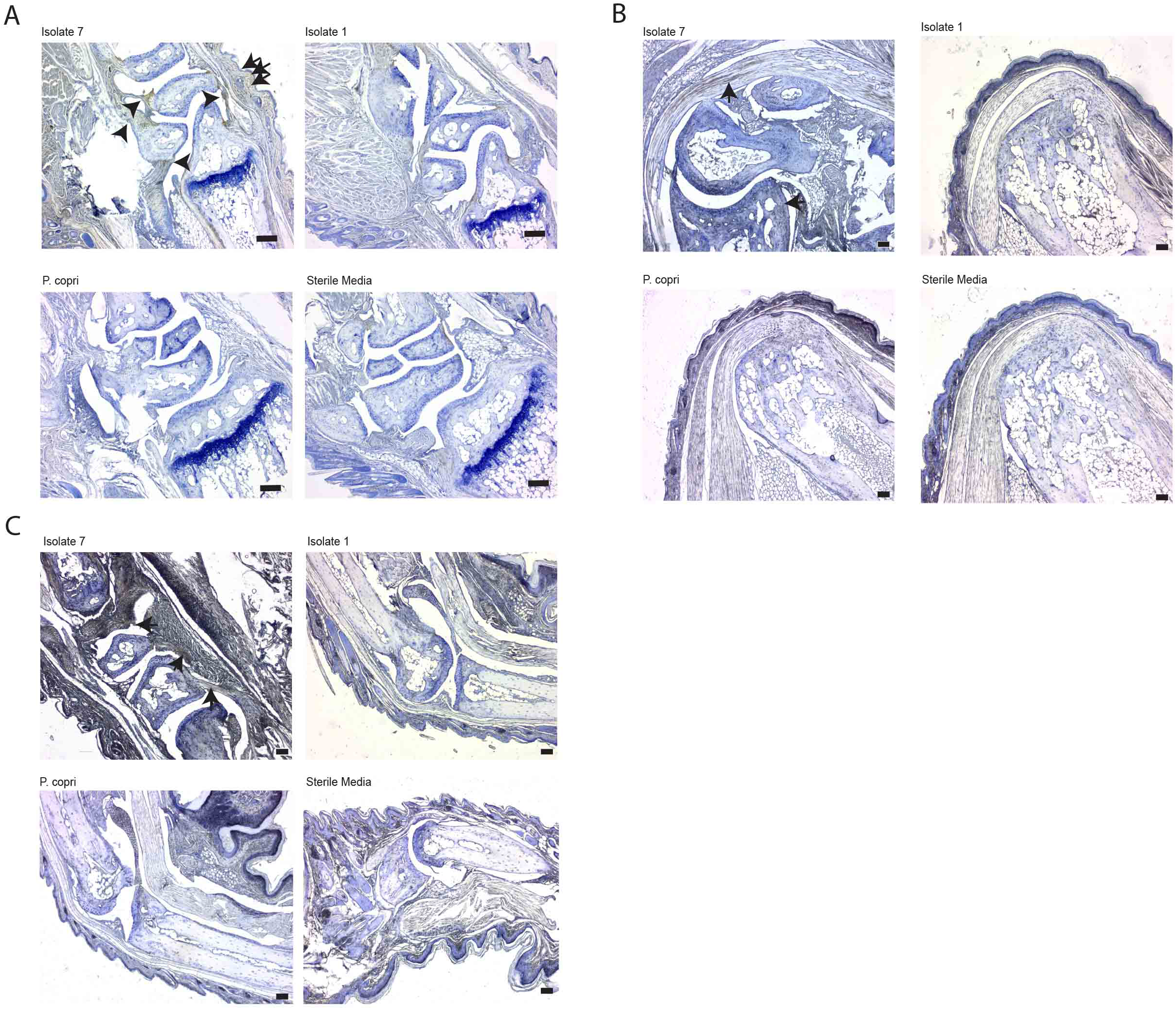
Complement C3, IgG, and IgA deposition in paws of mono-colonized and sterile media gavaged mice. Imrnunohistochemistry using antibodies targeting the C3 component of the complement cascade, IgG Fe or IgA Fe was performed on decalcified paw tissue sections. C3 deposition (A), IgG deposition (B), and IgA deposition (C) in representative sections is shown, with isolate 7 shown in the top left photo, isolate 1 shown in the top right photo, *P. copri* shown in the bottom left photo, and sterile media shown in the bottom right photo for each condition. Arrows denote regions of deposition, and scale bar is lOOum. (n=12 isolate 1 gavaged, n=11 isolate 7 gavaged, n=11 *P. copri* gavaged, and n=11 sterile media gavaged, across two experiments).

**Supplemental Figure 8:**
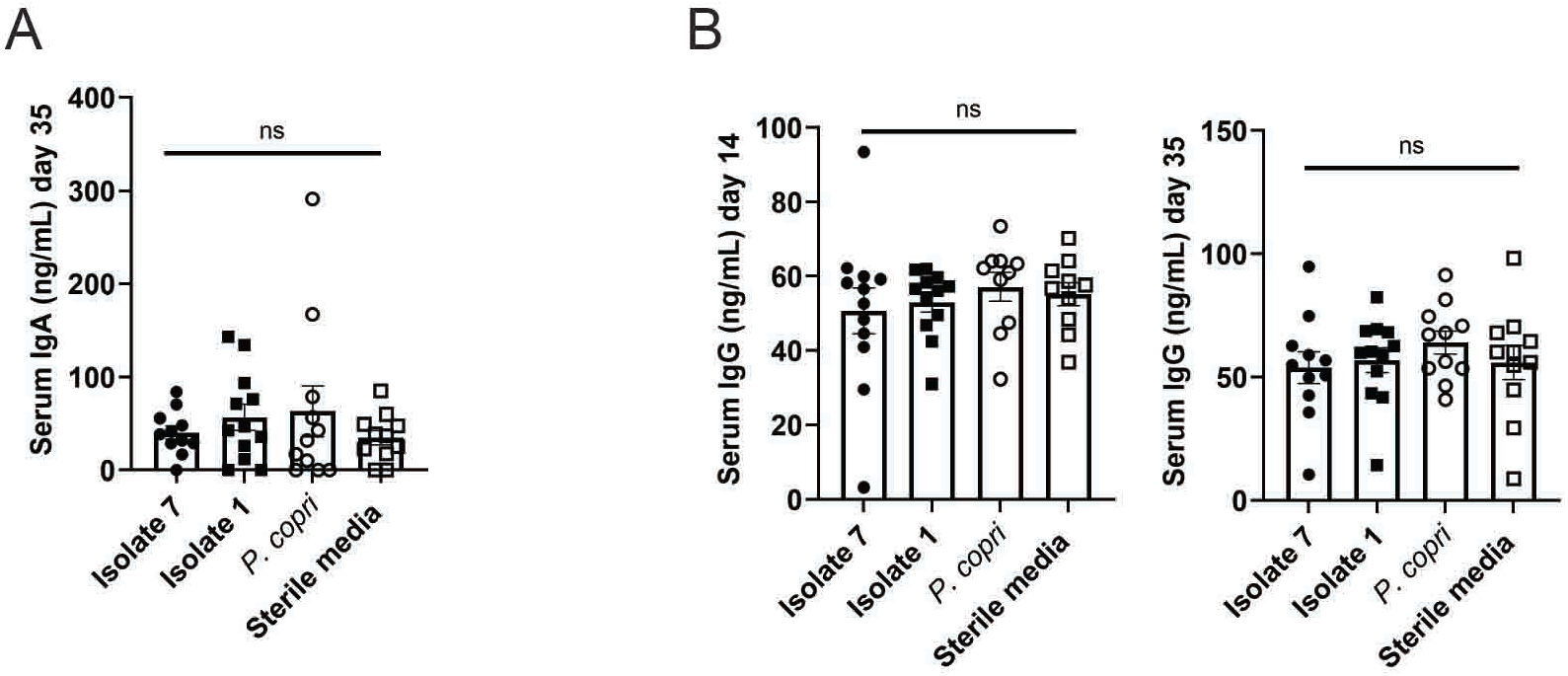
Serum IgA and IgG ofmonocolonized mice. Serum IgA (A) and IgG **(B)** levels were determined from the monocolonized mice at 14 days and 35 days after bacterial gavage. At day 14, n=l2 isolate gavaged mice, n=l2 isolate 1 gavaged mice, n=lO *P. copri* gavaged mice, and n=10 sterile media gavaged mice were tested. At day 35, n=11 isolate 7 gavaged mice, n=12 isolate 1 gavaged mice, n=11 *P. copri* gavaged mice, and n=11 sterile media gavaged mice were tested. The total IgA and IgG are displayed in ng/mL (y-axis), separated by treatment group (x-axis). ns= not significant as determined by Kruskal-Wallis test.

**Supplemental Figure 9:**
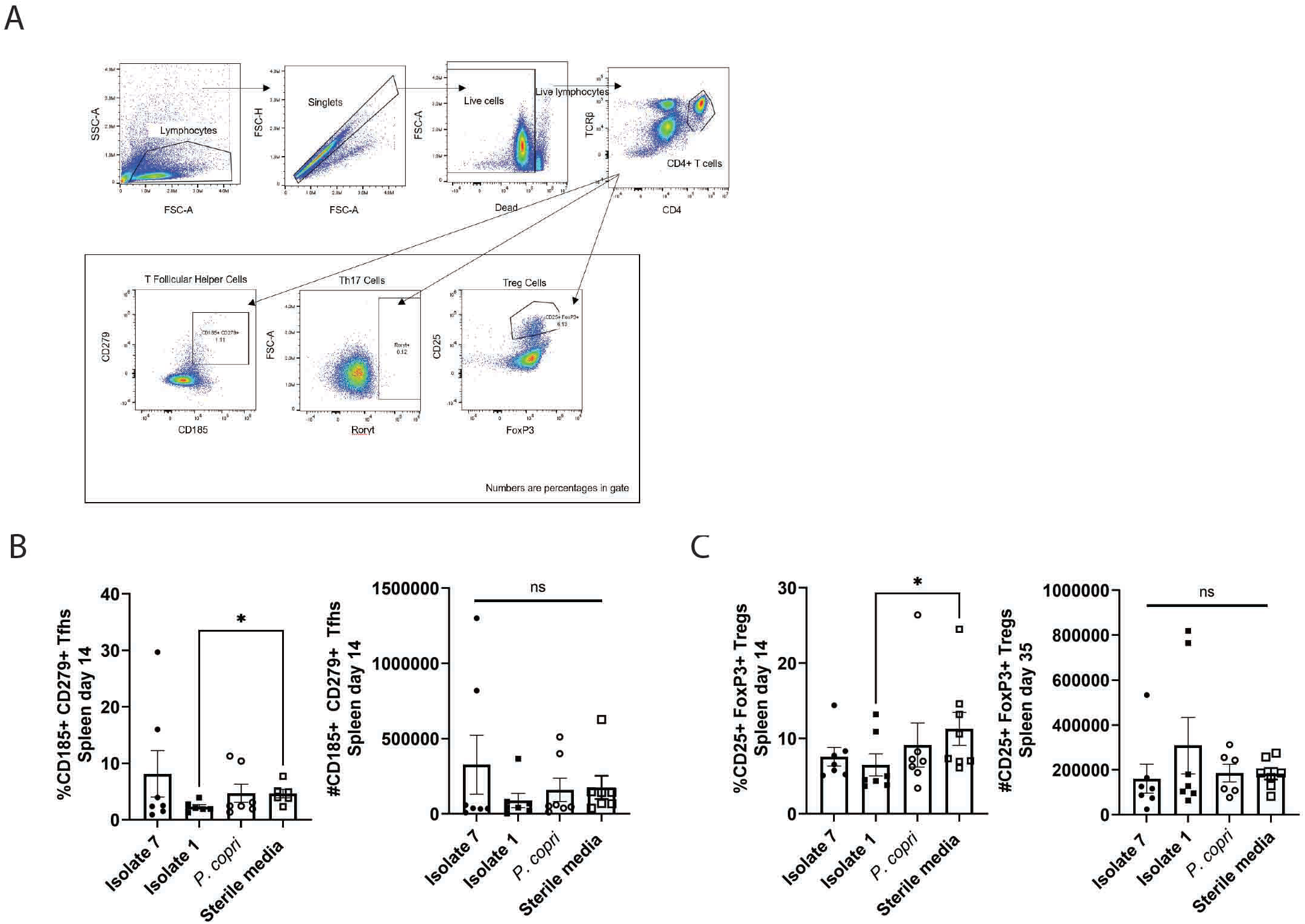
T cell immunophenotyping panel gating scheme. **(A)** The gating scheme for the T cell immunophenotyping panel is displayed here. Names of gates and percentages of cells in each gate are displayed. **(B)** Splenic T cell populations were determined at day 14 after gavage. The percentage and absolute number of T follicular helper cells (Live TCRp+ CD4+ CD185+ CD279+ lymphocytes) is displayed (y-axis) compared to treatment groups (x-axis). *P<0.05 and ns=not significant as determined by Kruskal-Wallis with Dunn’s post-test. (C) The percentage and absolute numbers ofTregs (Live TCRp+ CD4+ CD25+ FoxP3+ lymphocytes) are displayed (y-axis) compared to treatment group (x-axis). *P<0.05 and ns=not significant as determined by Kruskal-Wallis with Dunn’s post-test. n=7 isolate 7 gavaged mice, n=7 isolate 1 gavaged mice, n=7 *P. copri* gavaged mice, and n=8 sterile media gavaged mice.

**Supplemental Figure 10:**
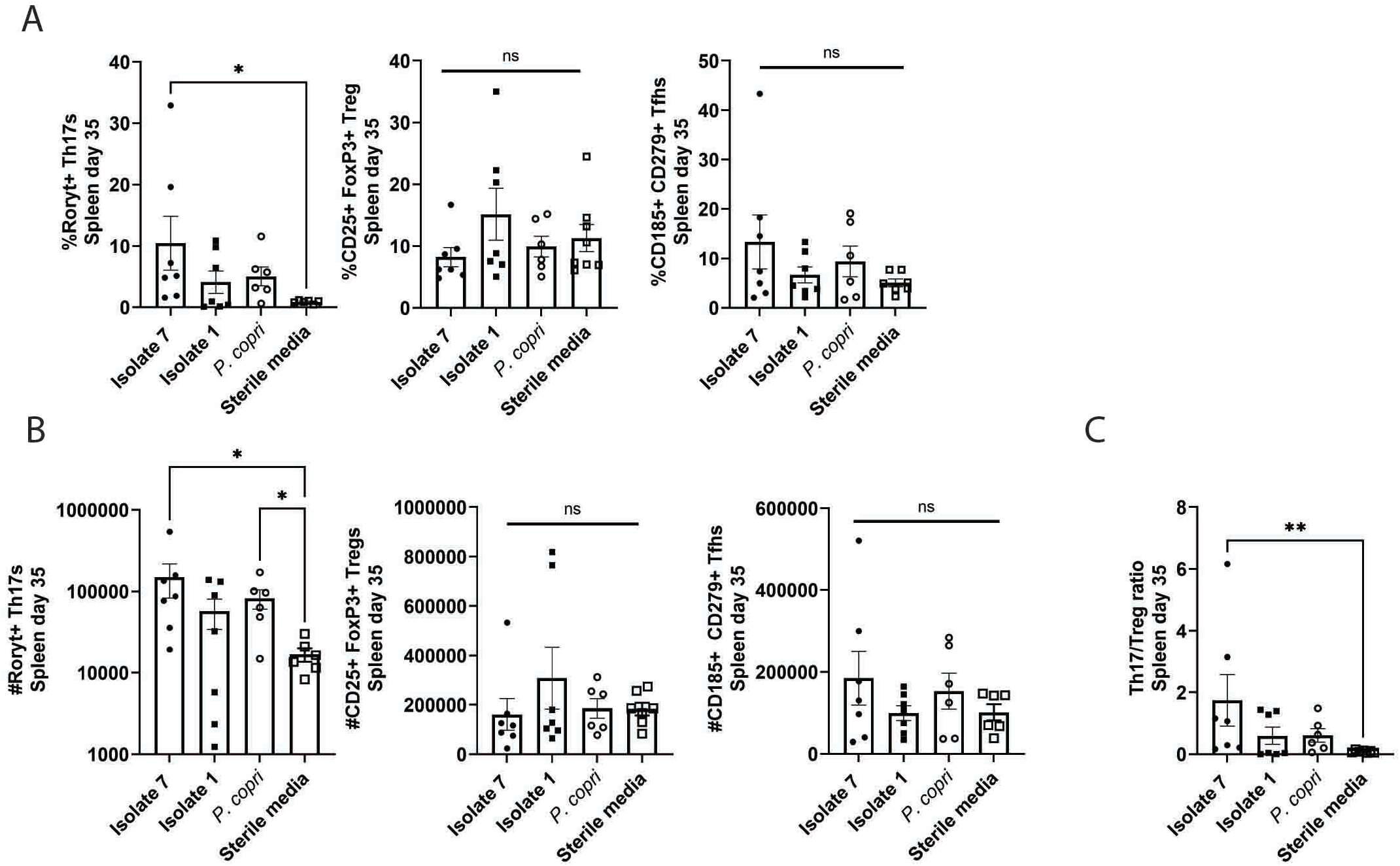
Splenic immune cell populations ofmonocolonized mice 35 days after bacterial gavage. Splenic T cell populations of treated mice were determined at day 35 after bacterial gavage. **(A)** The percentage ofThl 7s (Live TCRp+ CD4+ Roryt+ lymphocytes), Tregs (Live TCRp+ CD4+ CD25+ FoxP3+ lymphocytes), and T follicular helper cells (Live TCRp+ CD4+ CD185+ CD279+ lymphocytes) are displayed (y-axis), compared to treatment group (x-axis). (B) The absolute numbers of Thl 7s, Tregs, and Tfhs is displayed (y-axis) compared to treatment group (x-axis). (C) The ratio between Thl 7s and Tregs is shown (y-axis) compared to treatment group (x-axis) *P<0.05 and ns=not significant as determined by Kruskal-Wallis with Dunn’s post-test. n=7 isolate 7 gavaged mice, n=7 isolate 1 gavaged mice, n=6 *P. copri* gavaged mice, and n=8 sterile media gavaged mice.Supplemental Figure 11

**Supplemental Figure 11:**
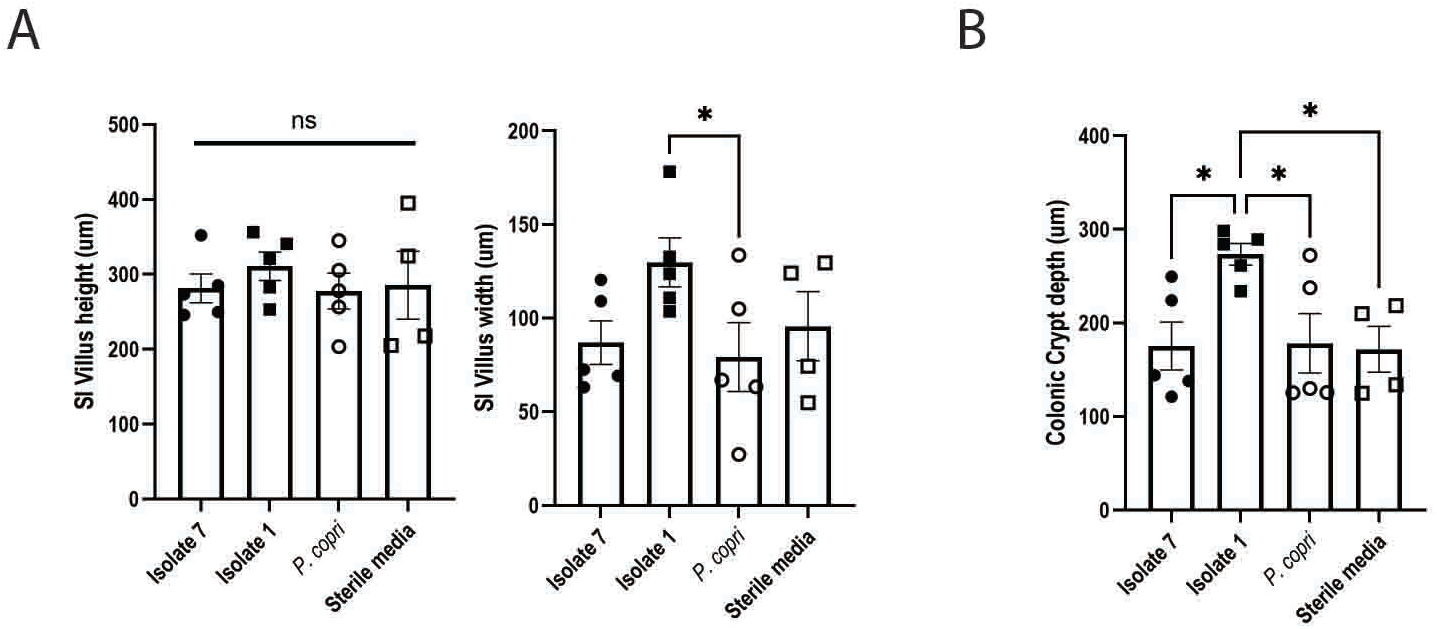
Small intestinal villus and crypt properties in monocolonized mice. **(A)** Small intestine villus height and width are displayed (y-axis, in microns) and separated by treatment group (x-axis), as determined by intestinal histology on gavaged mice. **(B)** Colon crypt depth (in microns) is displayed. *P<0.05 and ns=not significant as determined by one-way ANOVA with Tukey’s post-test. n= 5 mice per group.

**Supplemental Figure 12:**
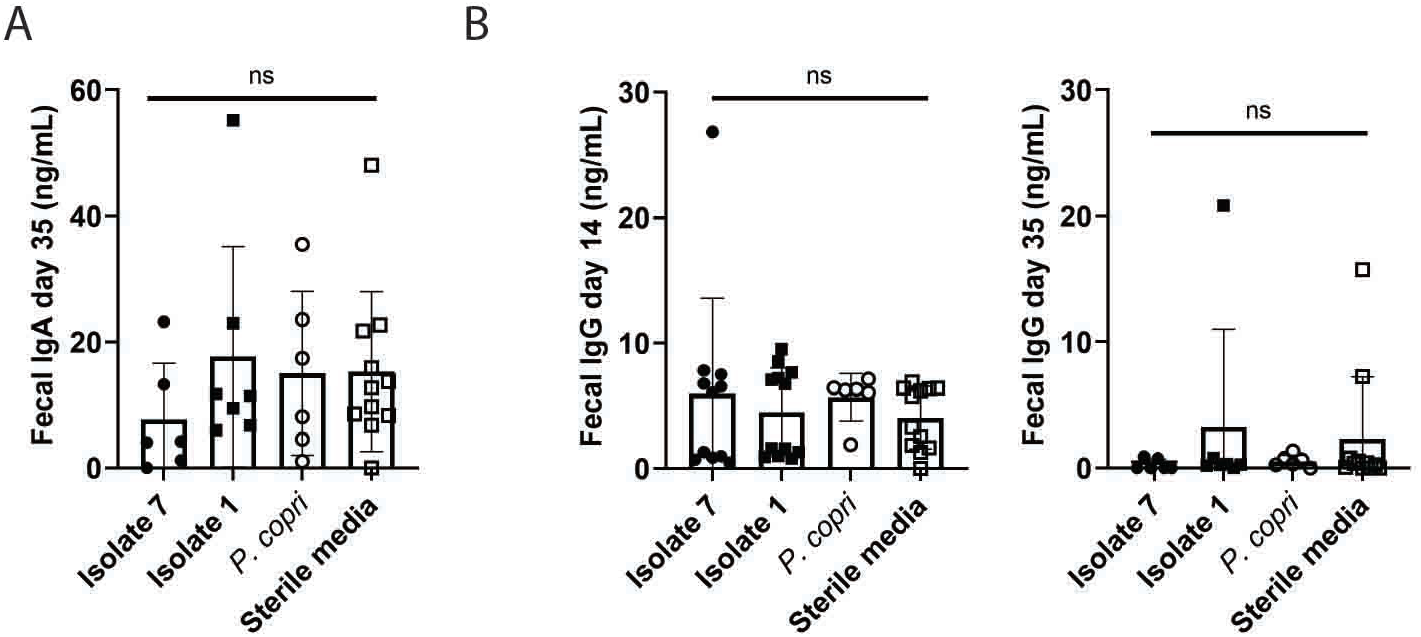
Fecal IgA and IgG of monocolonized mice. Fecal IgA **(A)** and IgG **(B)** levels were determined from the monocolonized mice at 14 days and 35 days after bacterial gavage. The total IgA and IgG are displayed in ng/mL, (y-axis) separated by treatment group (x-axis). For day 14, (n=l l isolate 7 gavaged mice, n=l2 isolate 1 gavaged mice, n=6 *P. copri* gavaged mice, and n=l2 sterile media gavaged mice were tested). For day 35, (n=6 isolate 7 gavaged mice, n=7 isolate 1 gavaged mice, n=6 *P. copri* gavaged mice, and n=11 sterile media gavaged mice were tested) (p-values determined by Kruskal-Wallis test).

**Supplemental Figure 13:**
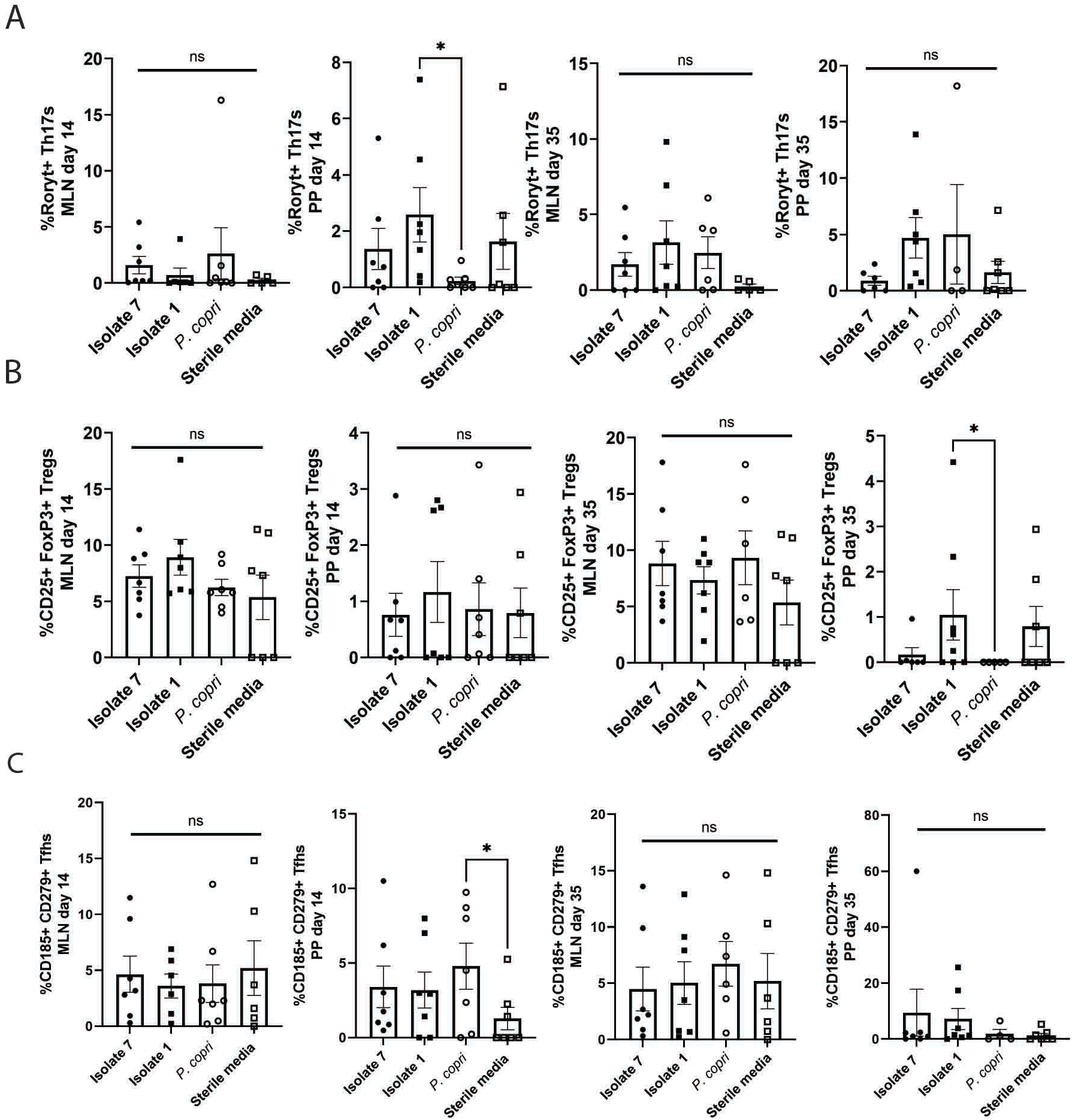
Mesenteric lymph node and Peyer’s patch immune cell populations in mice 14 and 35 days after gavage. MLN and Peyer’s patch T cells were determined at day 14 and 35 after gavage. **(A)** The percentage and absolute number of Thl 7s (live TCRp+ CD4+ Roryt+ lymphocytes) are shown (y-axis) compared to treatment group (x-axis). **(B)** The percentage and absolute number of Tregs (live TCRp+ CD4+ CD25+ FoxP3+ lymphocytes) are shown. (C) The percentage and absolute number of T follicular helper cells (live TCRp+ CD4+ CD185+ CD279+ lymphocytes) is shown. *P<0.05 and ns=not significant as determined by Kruskal-Wallis with Dunn’s post-test. n=7 isolate 7 gavaged mice, n=7 isolate l gavaged mice, n=7 *P. copri* gavaged mice, and n=8 sterile media gavaged mice.

**Supplemental Figure 14:**
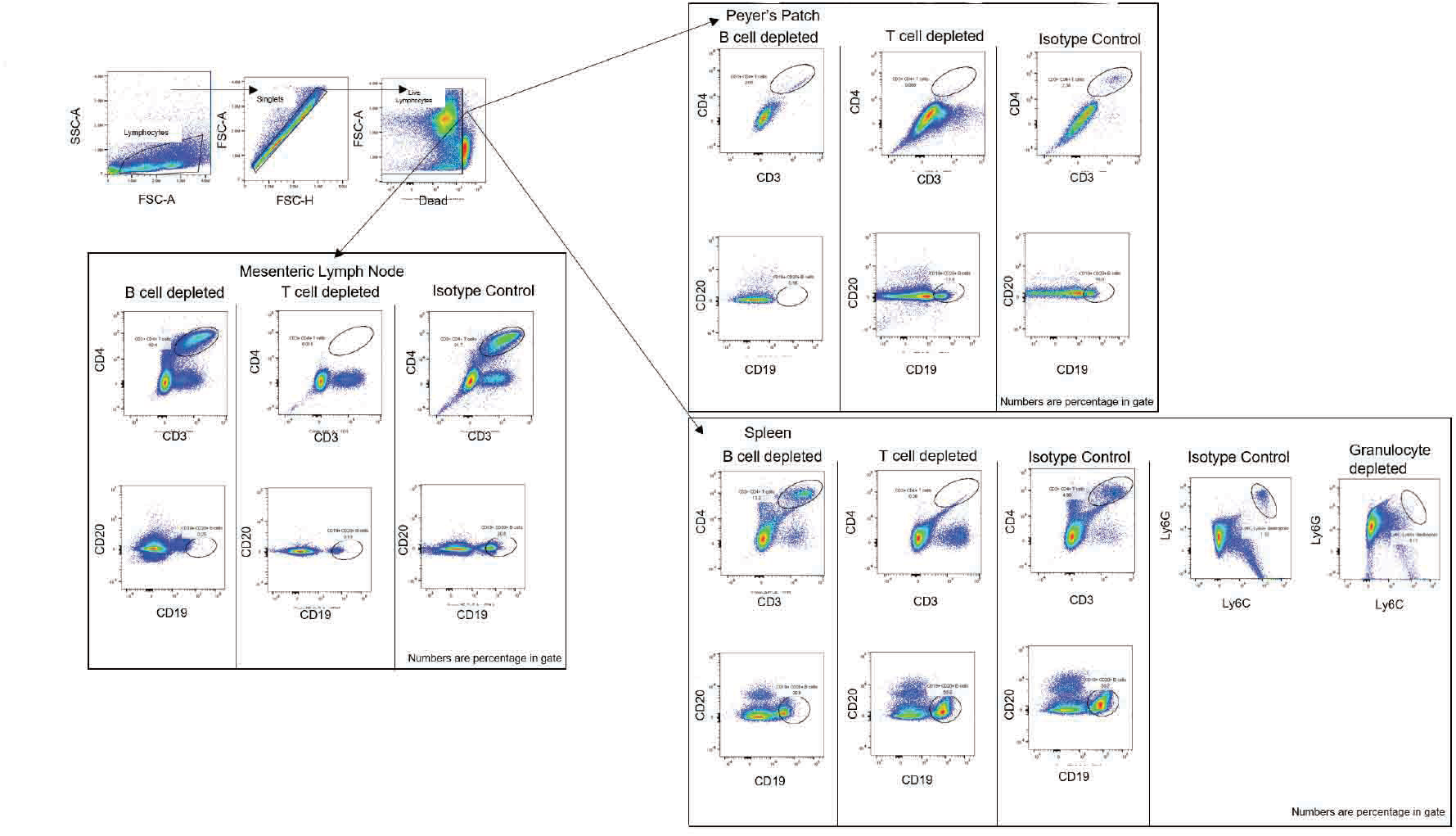
Gating scheme and validation of depletion antibodies. The gating scheme and validation of B cell, T cell, and granulocyte depletion antibodies is displayed for each tissue of interest, namely the MLNs, Peyer’s patches, and spleen. Each gate is labeled and contains the percentage of cells included.

**Supplemental Figure 15:**
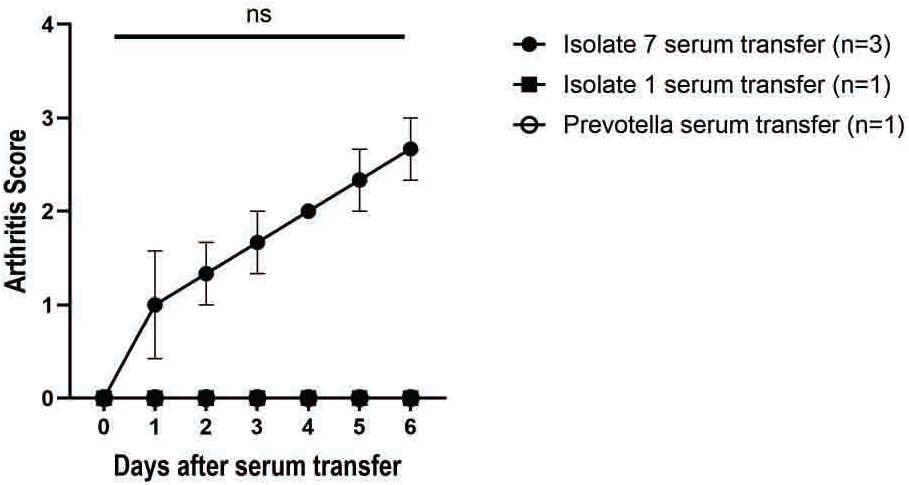
Mouse tissues collected at 6 days post-serum transfer before phenotype resolution occurs. Serum was collected from mice monocolonized with isolate **1,** isolate 7, and *P. copri* 35 days after bacterial gavage and was IP injected into healthy germ-free DBA/1 mice. These mice were monitored for the development of joint swelling and ankylosis every day for 6 days after serum transfer and then were euthanized for tissue collection. Clinical score is displayed (y-axis) against time since serum transfer (x­ axis) (p=0.002 at day 7, Mann-Whitney test) (n=4 isolate 7 serum transfer, n=4 isolate **1** serum transfer, n=4 *P. copri* serum transfer).

**Supplemental Table 1:** Characteristics of individuals in plasmablast mAb cohort.

| *Characteristic* | **At-risk subjects** | **eRA subjects** | **All subjects** |
| --- | --- | --- | --- |
| N | 4 | 2 | 6 |
| Sex: N (% female) | 3 (75.0) | 2 (100.0) | 5 (83.3) |
| Age: mean ± SD | 58.8 ± 8.7 | 44.5 ± 16.3 | 54.0 ± 12.3 |
| Race: n (% NHW) | 3 (75.0) | 1 (50.0) | 4 (66.7) |
| Ever Smoke: n (% yes) | 1 (25.0) | 2 (100.0) | 3 (50.0) |
| Current Smoker: n (% yes) | 0 (0.0) | 1 (50.0) | 1 (16.7) |
| Supplement Use: n (% yes) | 1 (25.0) | 0 (0.0) | 1 (16.7) |
| Other autoimmune disease: N (disease) |  |  |  |
| DMARD: n (% yes) |  |  |  |
| NSAID: n (% yes) | 1 (25.0) | 0 (0.0) | 1 (16.7) |
| Swollen Joint: n (% yes) | 0 (0.0) | 1 (50.0) | 1 (16.7) |
| Tender Joint: n (% yes) | 2 (50.0) | 1 (50.0) | 3 (50.0) |
| Tested for Shared Epitope: n (% positive) | 4 (100.0) | 1 (50.0) | 5 (83.3) |
| Shared Epitope: n (% positive *of those tested*) | 2 (50.0) | 1 (100.0) | 3 (60.0) |
| **CCP and RF Status** |  |  |  |
| RF- and CCP-: n (% yes) | 0 (0.0) | 0 (0.0) | 0 (0.0) |
| RF+ only: n (% yes) | 1 (25.0) | 0 (0.0) | 1 (16.7) |
| CCP+ only: n (% yes) | 1 (25.0) | 0 (0.0) | 1 (16.7) |
| RF+ and CCP+: n (% yes) | 2 (50.0) | 2 (100.0) | 4 (66.7) |
| **Type of CCP+** |  |  |  |
| Tested for CCP3: n (% yes) | 2 (50.0) | 1 (50.0) | 3 (50.0) |
| CCP3+: n (% yes *of those tested*) | 2 (100.0) | 1 (50.0) | 3 (50.0) |
| Tested for CCP3.1: n (% yes) | 4 (100.0) | 2 (100.0) | 6 (100.0) |
| CCP3.1+: n (% yes *of those tested*) | 3 (75.0) | 2 (100.0) | 5 (83.3) |
| **Type of RF+** |  |  |  |
| RF neph.+: n (% yes) |  |  |  |
| Tested for RF: n (% yes) | 4 (100.0) | 2 (100.0) | 6 (100.0) |
| RF IgA+: n (% yes *of those tested*) | 0 (0.0) | 2 (100.0) | 2 (33.3) |
| RF IgG+: n (% yes *of those tested*) | 1 (25.0) | 2 (100.0) | 3 (50.0) |
| RF IgM+: n (% yes *of those tested*) | 1 (25.0) | 2 (100.0) | 3 (50.0) |

Abbreviations: NHW = Non-Hispanic White; DMARD = disease modifying anti-rheumatic drug; NSAID

**Supplemental Table 2:** Characteristics of plasmablast cohort.

|  | **Subject 1** | **Subject 2** | **Subject 3** | **Subject 4** | **Subject 5** | **Subject 6** |
| --- | --- | --- | --- | --- | --- | --- |
| RA status | At-risk for RA development | At-risk for RA development | At-risk for RA development | At-risk for RA development | Early RA | Early RA |
| Number plasmablasts derived from subject | 18 | 18 | 18 | 24 | 7 | 9 |
| **Plasmablasts from each isotype in circulation** |  |  |  |  |  |  |
| %IgA+ plasmablasts | 61.14 | 55.19 | 49.65 | 68.94 | 26.40 | 0 |
| %IgG+ plasmablasts | 20.29 | 29.87 | 37.83 | 23.48 | 45.51 | 83.12 |
| %IgM+ plasmablasts | 4.29 | 7.14 | 5.44 | 2.27 | 11.8 | 12.52 |
| **Plasmablasts isolated for study (counts)** |  |  |  |  |  |  |
| Number of IgG plasmablasts | 6 | 2 | 6 | 11 | 5 | 8 |
| Number of IgA plasmablasts | 12 | 16 | 12 | 13 | 2 | 1 |
| **Plasmablasts targeting bacteria** |  |  |  |  |  |  |
| %Plasmablasts of cohort with bacterial targets | 66.67 | 83.33 | 56 | 71 | 43 | 11 |
| %IgG plasmablasts of cohort with bacterial targets | 67 | 100 | 67 | 64 | 60 | 13 |
| %IgA plasmablasts of cohort with bacterial targets | 67 | 81 | 50 | 77 | 0 | 0 |

**Supplemental Table 4:** Characteristics of human samples used for fecal pool.

| Characteristic | Healthy Controls | At-risk | Early RA |
| --- | --- | --- | --- |
| N | 5 | 8 | 5 |
| Sex: N (% female) | 4 (80) | 6 (75) | 3 (60) |
| Age: mean ± SD | 41.2 ± 13.6 | 62 ± 11.3 | 46.2 ± 14.5 |
| Race: N (% NHW) | 4 (80) | 7 (87.5) | 4 (80) |
| Ever smoke: N (% yes) | 1 (20) | 1 (12.5) | 1 (20) |
| Current smoker: N (%yes) | 0 (0) | 0 (0) | 0 (0) |
| Supplement use: N (%yes) | 4 (80) | 6 (75) | 5 (100) |
| DMARD: N (% yes) | 0 (0) | 0 (0) | 4 (80) |
| NSAID: N (% yes) | 1 (20) | 5 (62.5) | 2 (40) |
| Swollen joint: N (% yes) | 0 (0) | 1 (12.5) | 3 (60) |
| Tender joint: N (% yes) | 1 (20) | 4 (50) | 3 (60) |
| Shared Epitope: N (% positive) | 4 (80) | 3 (37.5) | 3 (60) |
| **CCP and RF status** |  |  |  |
| RF- and CCP-: N (% yes) | 5 (100) | 2 (25) | 1 (20*) |
| RF+ only: N (% yes) | 0 (0) | 1 (12.5) | 1 (20) |
| CCP+ only: N (% yes) | 0 (0) | 1 (12.5) | 1 (20) |
| RF+ and CCP+: N (% yes) | 0 (0) | 4 (50) | 2 (40) |
| **Type of CCP** |  |  |  |
| CCP3+: N (% positive) | 0 (0) | 3 (37.5) | 3 (60) |
| CCP3.1+: N (% positive) | 0 (0) | 5 (62.5) | 3 (60) |
| **Type of RF** |  |  |  |
| RF neph+: N (% yes, *of those tested*) | 0 (0) | 2 (28.6) | 0 (0) |
| RF IgA+: N (% positive) | 0 (0) | 1 (12.5) | 2 (40) |
| RF IgG+: N (% positive) | 0 (0) | 2 (25) | 0 (0) |
| RF IgM+: N(% positive) | 0 (0) | 4 (50) | 2 (40) |

Abbreviations: NHW = Non-Hispanic White; DMARD = disease modifying anti-rheumatic drug; NSAID = non-steroidal anti-inflammatory drug; CCP = cyclic citrullinated peptide; RF = rheumatoid factor. \* This individual was previously CCP2+, CCP3.1+, and IgG RF+ but had negative titers at the time of draw.

**Supplemental Table 5:** Characteristics of RA cases for CD4+ T cell reactivity assays.

| *Characteristic* | **At-risk**<br>**subjects** | **eRA**<br>**subjects** | **All subjects** |
| --- | --- | --- | --- |
| N | 4 | 2 | 6 |
| Sex: N (% female) | 3 (75.0) | 2 (100.0) | 5 (83.3) |
| Age: mean ± SD | 58.8 ± 8.7 | 44.5 ± 16.3 | 54.0 ± 12.3 |
| Race: n (% NHW) | 3 (75.0) | 1 (50.0) | 4 (66.7) |
| Ever Smoke: n (% yes) | 1 (25.0) | 2 (100.0) | 3 (50.0) |
| Current Smoker: n (% yes) | 0 (0.0) | 1 (50.0) | 1 (16.7) |
| Supplement Use: n (% yes) | 1 (25.0) | 0 (0.0) | 1 (16.7) |
| Other autoimmune disease: N (disease) |  |  |  |
| DMARD: n (% yes) |  |  |  |
| NSAID: n (% yes) | 1 (25.0) | 0 (0.0) | 1 (16.7) |
| Swollen Joint: n (% yes) | 0 (0.0) | 1 (50.0) | 1 (16.7) |
| Tender Joint: n (% yes) | 2 (50.0) | 1 (50.0) | 3 (50.0) |
| Tested for Shared Epitope: n (% positive) | 4 (100.0) | 1 (50.0) | 5 (83.3) |
| Shared Epitope: n (% positive *of those tested*) | 2 (50.0) | 1 (100.0) | 3 (60.0) |
| **CCP\* and RF† Status** |  |  |  |
| RF- and CCP-: n (% yes) | 0 (0.0) | 0 (0.0) | 0 (0.0) |
| RF+ only: n (% yes) | 1 (25.0) | 0 (0.0) | 1 (16.7) |
| CCP+ only: n (% yes) | 1 (25.0) | 0 (0.0) | 1 (16.7) |
| RF+ and CCP+: n (% yes) | 2 (50.0) | 2 (100.0) | 4 (66.7) |
| **Type of CCP+** |  |  |  |
| Tested for CCP3: n (% yes) | 2 (50.0) | 1 (50.0) | 3 (50.0) |
| CCP3+: n (% yes *of those tested*) | 2 (100.0) | 1 (50.0) | 3 (50.0) |
| Tested for CCP3.1: n (% yes) | 4 (100.0) | 2 (100.0) | 6 (100.0) |
| CCP3.1+: n (% yes *of those tested*) | 3 (75.0) | 2 (100.0) | 5 (83.3) |
| **Type of RF+** |  |  |  |
| RF neph.+: n (% yes) |  |  |  |
| Tested for RF: n (% yes) | 4 (100.0) | 2 (100.0) | 6 (100.0) |
| RF IgA+: n (% yes *of those tested*) | 0 (0.0) | 2 (100.0) | 2 (33.3) |
| RF IgG+: n (% yes *of those tested*) | 1 (25.0) | 2 (100.0) | 3 (50.0) |
| RF IgM+: n (% yes *of those tested*) | 1 (25.0) | 2 (100.0) | 3 (50.0) |

**Supplemental Table 6:** Characteristics of RA and Healthy Control cases for CD4+ T cell reactivity assays.

| Event ID | Patient Type | Anti-CCP Outcome | Rheumatoid Factor Outcome | HLA DRB1(a+b) | Gender | Age at Draw | Rheumatoid Arthritis: Duration at Draw | Active Autoimmune Medications |
| --- | --- | --- | --- | --- | --- | --- | --- | --- |
| Ce1381922 | HC | Negative | Negative | *0401/*1501 | Female | 57 | N/A | None |
| Ce1265824 | HC | Negative | Negative | *0301/*0401 | Male | 29 | N/A | None |
| Ce1359061 | HC | Negative | Negative | *1402/*0401 | Female | 39 | N/A | None |
| Ce1594449 | HC | Negative | Negative | *0401/*0405 | Female | 40 | N/A | None |
| Ce1352794 | RA | Positive | Positive | *0101/*0401 | Female | 73 | 0 | Methotrexate |
| Ce1352794 | RA | Positive | Positive | *0405/Unknown | Female | 19 | 3.4 | Etanercept |
| Ce913294 | RA | Positive | Positive | *0102/*11 | Female | 72 | 0.6 | Meloxicam, Infliximab |
| Ce1371765 | RA | Positive | Positive | *01/*0401 | Female | 20 | 2.9 | Tofacitinib |

Abbreviations: CCP = cyclic citrullinated peptide

**Supplemental Table 8:** T cell immunophenotyping panel information.

| Antibody Target | Fluorophore | Clone | Company |
| --- | --- | --- | --- |
| anti-mouse CD3 | PerCP Cy5.5 | 145-2C11 | Tonbo |
| anti-mouse CD4 | PE | GK1.5 | Tonbo |
| anti-mouse CD25 | FITC | PC61.5 | Tonbo |
| anti-mouse CD185 | APC-Cy7 | L138D7 | Biolegend |
| anti-mouse CD279 | BV421 | 29F.1A12 | Biolegend |
| anti-mouse Foxp3 | PE-Cy7 | 3G3 | Tonbo |
| anti-mouse Roryt | BV786 | Q31-378 | BD Horizon |
| anti-mouse Tbet | PE Vio615 | REA102 | Miltenyi |
| anti-mouse TCRβ | APC | H57-597 | Tonbo |
| viability | Ghost510 | n/a | Tonbo |

**Supplemental Table 9:**
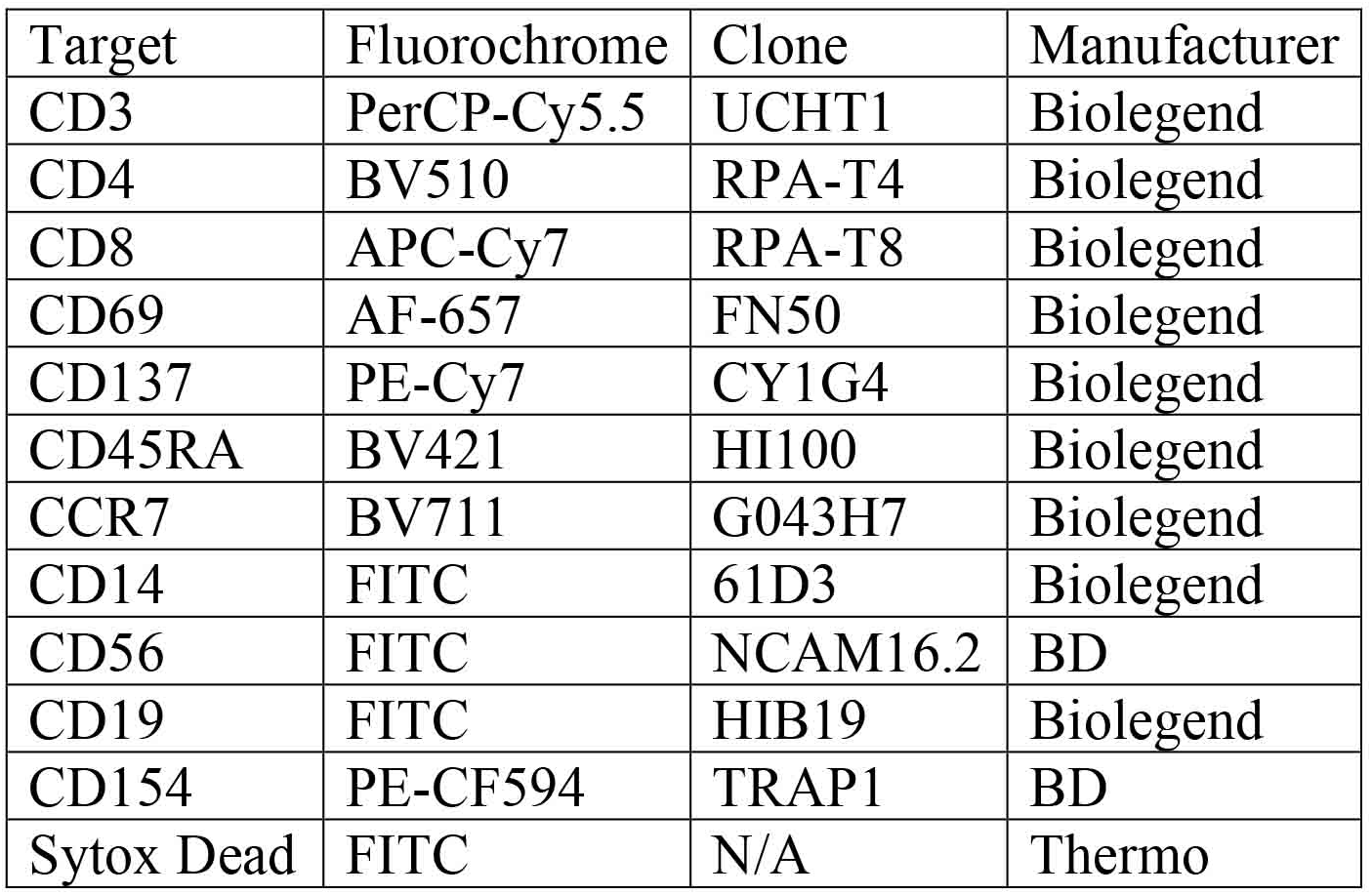
Human T cell stimulation antibody information.

| Target | Fluorochrome | Clone | Manufacturer |
| --- | --- | --- | --- |
| CD3 | PerCP-Cy5.5 | UCHT1 | Biolegend |
| CD4 | BV510 | RPA-T4 | Biolegend |
| CD8 | APC-Cy7 | RPA-T8 | Biolegend |
| CD69 | AF-657 | FN50 | Biolegend |
| CD137 | PE-Cy7 | CY1G4 | Biolegend |
| CD45RA | BV421 | HI100 | Biolegend |
| CCR7 | BV711 | G043H7 | Biolegend |
| CD14 | FITC | 61D3 | Biolegend |
| CD56 | FITC | NCAM16.2 | BD |
| CD19 | FITC | HIB19 | Biolegend |
| CD154 | PE-CF594 | TRAP1 | BD |
| Sytox Dead | FITC | N/A | Thermo |

## Notes

### Competing Interest Statement

The authors have declared no competing interest.

